# Shared molecular consequences of epigenetic machinery disruption in murine neuronal progenitors

**DOI:** 10.1101/2025.09.12.675897

**Authors:** Juan Ouyang, Kaan Okay, Katrín Möller, Katrín Wang, Adam Davidovich, Stefán Pétursson, Tessa Pierce, Arsalan Amirfallah, Kimberley J. Anderson, Kasper D. Hansen, Hans Tomas Bjornsson

## Abstract

The Mendelian disorders of the epigenetic machinery (MDEMs) are an emerging cause of intellectual disability and growth abnormalities, which commonly disrupt hippocampal function. To investigate consequences of epigenetic machinery (EM) disruption during neurodevelopment, we systematically knocked out (KO) EM factors in neuronal progenitors isolated from the murine hippocampus and established a neurodifferentiation model to interrogate their functions. We then profiled gene expression and DNA methylation (DNAm) in the EM-KOs using RNA sequencing and Nanopore long-read DNA sequencing. While *Dnmt1-*KO induced extensive DNAm alterations, *Kmt2a*-KO had little effect on methylation. Nevertheless, the disruption of *Kmt2a* and *Dnmt1* produced strikingly convergent transcriptional changes. Loss of either EM factor led to premature neuronal differentiation, partially explaining this convergence, and MYC appeared as a shared regulatory node linked to downregulation of cell-cycle programs in these cells. Extending our methylation analysis to 46 EM genes, we found that loss of DNA methyltransferases induced the strongest DNAm changes, whereas other EM-KOs had subtle or negligible effects. However, clustering of EM-KOs based on promoter DNAm levels revealed three distinct EM subgroups, of which two were enriched for interactions with the DNAm machinery. Allele-specific analysis of DNAm further identified a single differentially methylated region shared across the 46 EM-KOs, localized to the FVB allele over the *Zic4* 3’UTR. Furthermore, *Zic4* overexpression appears to maintain the neuronal progenitor state, suggesting functional relevance of this locus. Taken together, our results reveal both gene-specific and convergent effects across diverse EM-KOs and provide new insight into the molecular etiology of the MDEMs.

## Introduction

In the past 15 years clinicians have witnessed the rapid discovery of a group of at least 90 distinct disorders, caused mainly by heterozygous variants in the components that directly maintain the epigenome. This group of disorders is described by different names, including the Mendelian disorders of the epigenetic machinery (MDEMs) (Bjornsson, 2015), chromatinopathies (Ciptasari & van Bokhoven, 2020), or rare diseases of epigenetic origin (Fu et al., 2023). Despite this multitude of names, all parties agree that the disruption of these genes is likely to lead to secondary epigenetic dysregulation. Although these genes have been associated with broad phenotypic presentations (Bjornsson, 2015), they also exhibit unifying disease phenotypes such as intellectual disability and growth abnormalities (Bjornsson, 2015) which are present in the vast majority of patients. We have previously hypothesized that these shared phenotypic abnormalities may be caused by shared epigenetic disruption. In support of this hypothesis, we have identified common signatures of epigenetic dysregulation in three such MDEMs (Kabuki syndrome 1&2 and Rubinstein-Taybi syndromes): identified in blood cells for all three disorders (Luperchio et al., 2021) and in neurons for the first two (Boukas et al., 2024).

Another defining feature of these disorders is that many of them (42) exhibit distinct blood cell DNA methylation (DNAm) episignatures with diagnostic utility (Aref-Eshghi et al., 2020). Remarkably, although blood may not always be the most disease-relevant tissue, many of these disorders have shared epigenetic abnormalities (Aref-Eshghi et al., 2018). Here, we have investigated whether such DNAm changes can be found in disease-relevant cells, such as neurons, and whether they can point to underlying shared pathogenic mechanisms in these disorders. To do so, we have focused on a hippocampal neuronal model, given prior evidence of disruption of hippocampal neurogenesis in many of the MDEM (Carosso et al., 2019; Feng et al., 2013; Noguchi et al., 2016; Park et al., 2014; Schoof et al., 2019; Sheu et al., 2019; Syal et al., 2018; You et al., 2015; Zhang et al., 2014; Zhou et al., 2016).

Recent advances in Oxford Nanopore Technologies (ONT) long-read direct DNA sequencing allow for the analysis of DNAm genome-wide and further enables haplotype phasing for the identification of allele-specific effects on DNAm (Gigante et al., 2019). Here, we profiled DNAm abnormalities seen in 46 EM knockouts (KOs) in a single disease-relevant cell-type context, murine neuronal progenitor cells (mNPCs). Our work identifies several transcription factors (TFs) which appear to mediate aspects of the observed chromatin abnormalities or show dysregulation secondarily to the epigenetic abnormality. Our approach, which enables haplotype phasing of the DNAm data, also highlights the considerable effect that strain-specific genetic variation has on DNAm levels. Together, our results lay the groundwork for understanding how the epigenetic machinery (EM) helps orchestrate hippocampal neurogenesis and how disruption of this process through EM loss uniformly leads to learning and neurological defects.

## Results

### Systematic KO of EM factors in a single neuronal system

To comprehensively study the role of DNAm abnormalities in EM-deficient neurons, we utilized CRISPR-Cas9 technology to generate homozygous KO of EM genes in mNPCs. To obtain homogenous and efficient CRISPR-based gene KOs, we used F1 offspring from two fully sequenced inbred mouse-strains (C57BL/6J and FVB), both expressing transgenic Cas9-GFP protein from the ROSA locus, a well-known epigenetic safe harbor region (Friedrich & Soriano, 1991; Irion et al., 2007). We isolated hippocampal mNPCs at postnatal day 0 (P0) from this F1-outcross and grew them in culture media that maintains the mNPC state. We verified cell identities (**Supplementary Figure 1A**) prior to individual EM-KO experiments using 46 dual guide-RNA (sgRNA) lentiviral plasmids (**Supplementary Table 1**), each containing two sgRNA sequences targeting the same gene. We also created eight sgRNA plasmids with random sequences (scramble gRNAs, **Supplementary Table 1**) for use as controls. These guide-vectors were individually packaged and transduced using lentiviral delivery into the Cas9-expressing mNPCs (**Figure 1A**). Cells were selected for stable integration of a construct and, four days after transduction, harvested for profiling of DNAm (ONT sequencing) or gene expression (Illumina RNA-seq).

**Figure 1:**
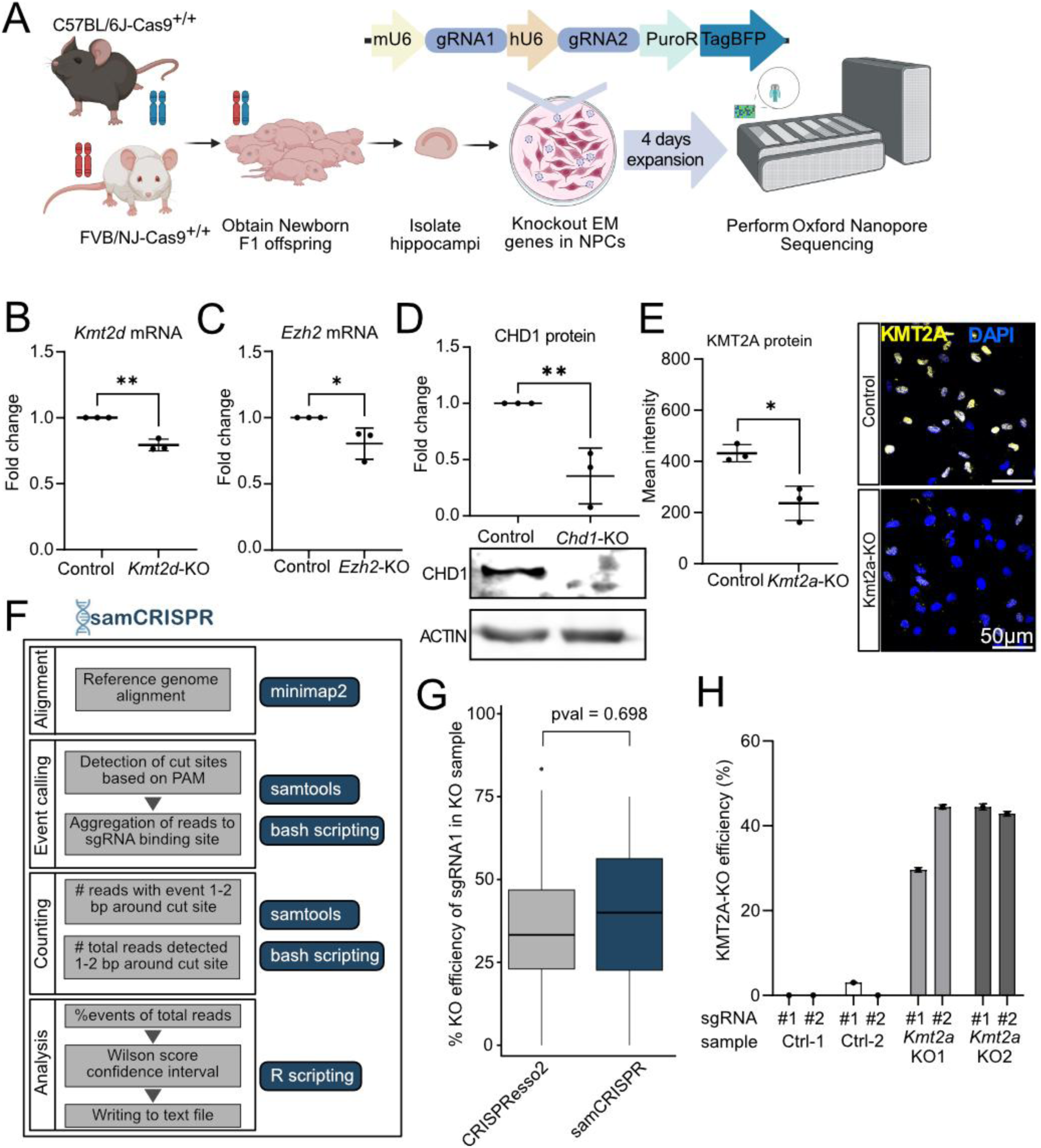
Effective CRISPR-Cas9 targeting in a disease relevant cellular system. (A) Schematic workflow of EM-KO in mNPCs using lentiviral vectors followed by evaluation of DNAm with ONT sequencing. (B) Gene expression changes of *Kmt2d* (p=0.0013) and (C) *Ezh2* (p=0.0229) in mNPCs after knocking out *Kmt2a* or *Ezh2* with CRISPR/Cas9 compared with scrambled control (n=3). An unpaired t-test was used to calculate significance. (D) Protein levels and quantification of CHD1 after knocking out *Chd1* with CRISPR-Cas9, compared with scrambled control (n=3, p=0.0054). An unpaired t-test was used to calculate significance. (E) Quantification of *Kmt2a* mean intensity per cell in *Kmt2a*-KO and scrambled controls (n=3, p=0.0001). An unpaired t-test was used to calculate significance. (F) Schematic of the *samCRISPR* analysis tool for KO efficiency estimation of long-read sequencing data. (G) The KO efficiency estimation by samCRISPR method vs CRISPResso2, for individual sgRNAs in 46 EM-KO from ONT WGS sequencing data (Two tailed t-test). (H) Bar plot showing *Kmt2a-*KOs efficiencies for sgRNA #1 and #2 in two *Kmt2a*-KOs and controls (Ctrl) calculated by samCRISPR. *p<0.05, **p<0.01, ***p<0.001, ns=not significant.

To confirm the efficiency of the KO in our system, we performed qPCR, western blot (WB), and immunofluorescence (IF) assays on individual EM-KO mNPCs. As expected, four days after transduction, we routinely saw a decrease in the relevant mRNA and protein levels for individual EM-KOs. As an example, we observed significant (*p* < 0.05) downregulation of *Kmt2d* and *Ezh2* at the RNA level (**Figure 1B-C**) as well as a significant (*p* < 0.05) decrease in protein levels of CHD1, KMT2A, and DNMT1 (**Figure 1D-E, Supplementary Figure 1B-C**) in each of their respective KOs.

To enable accurate and systematic estimation of CRISPR KO efficiency from ONT long-read sequencing data, we developed *samCRISPR*, a custom pipeline designed to leverage the unique features of this sequencing technique. This approach utilizes Minimap2 (Li, 2018) for sequence alignment and SAMtools (Danecek et al., 2021) for downstream processing (**Figure 1F**). When we compared *samCRISPR* with *CRISPResso2* (Pinello et al., 2016), the gold standard for estimating KO efficiency from short-read data, it yielded fewer false positive CRISPR events on average in control samples (2.1% vs 4.9% for sgRNA1, p < 0.001, and 1.4% vs 4.5% for sgRNA2, p < 2.2e-16)); **Supplementary Figure 1D-E**). Moreover, *samCRISPR* yielded a similar KO efficiency for each guide (39.1% vs 37.5% for sgRNA1 and 36.85% vs 35.3% for sgRNA2; **Figure 1G, Supplementary Figure 1F-G**) across 46 EM-KOs. As an example, when we ran two replicates of *Kmt2a*-KO and two relevant controls, we found almost no false positives in the control samples, while we detected on average 40% KO efficiency for either allele of the *Kmt2a*-KOs (**Figure 1H**). An underestimate of the actual KO efficiency is expected for both methods due to the small detection window (+/- 1bp around cut-site), and both WB and IF showed higher KO efficiency, i.e. lower protein expression (40-90%) in *Chd1-*, *Kmt2d-*, and *Dnmt1-*KOs (**Figure 1D-E, Supplementary Figure 1B-C**) supporting the notion that a portion of the functional variants are not captured by either samCRISPR or CRISPResso2.

### DNAm and transcriptional changes in *Dnmt1*-KO and *Kmt2a*-KO mNPCs

Having verified the KO efficiency of our system, we investigated if the disruption of individual EM genes in our mNPCs would result in DNAm abnormalities. As *Dnmt1* encodes the key maintenance DNA methyltransferase, we used it as a positive control for assessing the ONT sequencing approach. We performed ONT sequencing (Jain et al., 2016) on *Dnmt1-*KO and scrambled controls in biological triplicates. Differential methylation analysis revealed a global demethylation pattern in *Dnmt1*-KO cells, including 141,721 hypomethylated differentially methylated regions (DMRs) (qval < 0.05, **Figure 2A, Supplementary Table 2**) and a sole hypermethylated region in *Zic1* gene body (**Figure 2B**). This is consistent with prior studies showing that loss of DNMT1 in human NPCs causes global demethylation (Jonsson et al., 2019). *Dnmt1-*KO DMRs were enriched in 3’UTRs, introns, and exons but depleted in promoters, intergenic regions, and 5’UTRs (**Figure 2C, Supplementary Table 3**).

**Figure 2:**
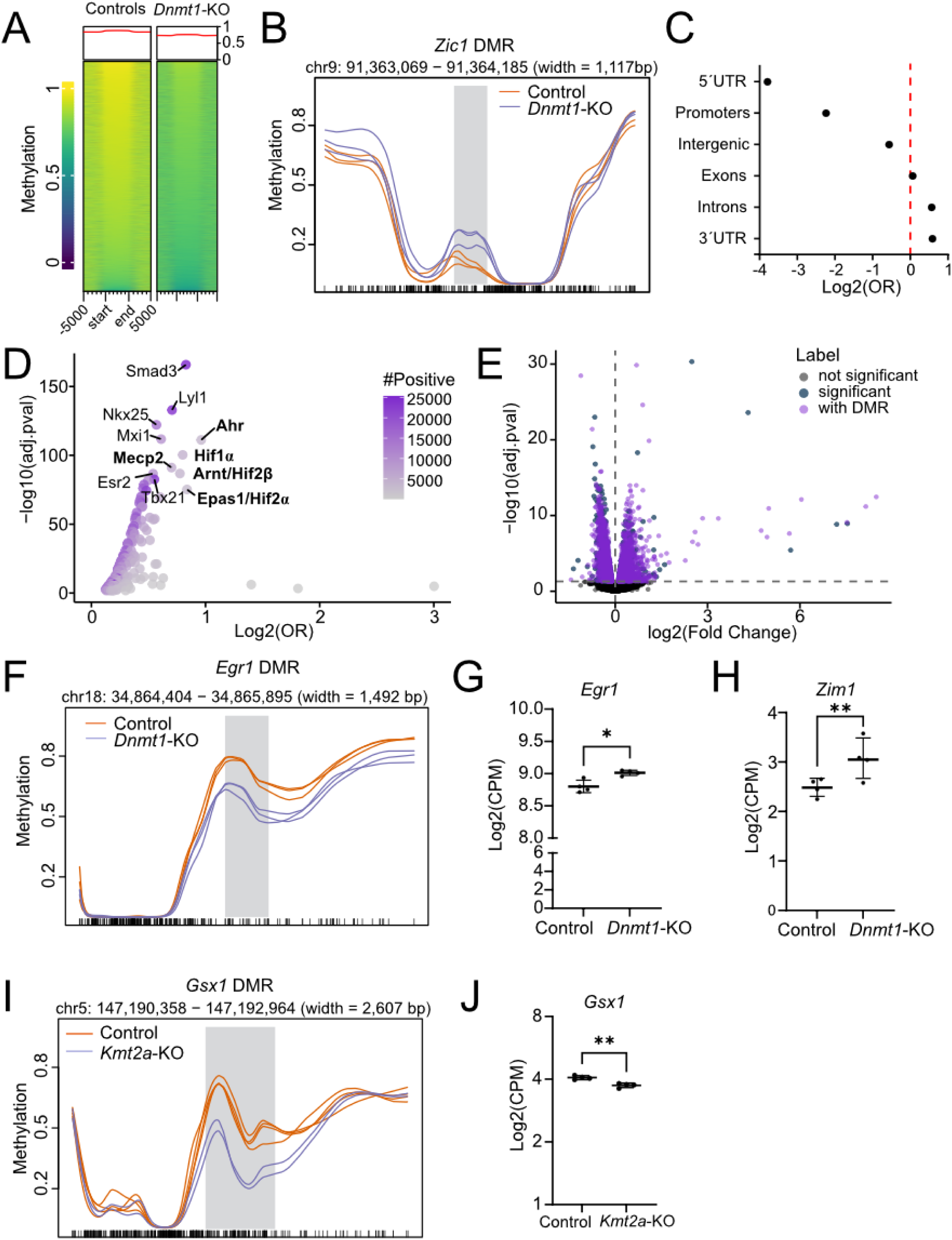
Extent and impact of DNA methylation changes in *Dnmt1*-KO and *Kmt2a*-KO mNPCs. (A) Enrichment heatmaps of smoothed CpG methylation across 141,722 *Dnmt1*-associated DMRs. Each horizontal line represents the average smoothed methylation profile across CpG sites within a DMR and ±5kb flanking regions. Yellow shows higher smoothed methylation but blue shows lower smoothed methylation. (B) The *Zic1* DMR (qval < 0.05, gray vertical bar) in *Dnmt1*-KOs (purple) compared to control NPCs (orange). Flanking regions of DMR are ±5kb. On the x-axis, each tick mark represents a CpG in the mouse genome. (C) The enrichment analysis of the DMR distribution in *Dnmt1*-KOs, using Fisher’s exact test. All categories are significantly enriched or depleted (p < 0.05). The exact value is listed in **Supplementary Table 3**. (D) Scatter plot showing enriched TF motifs in *Dnmt1*-KO DMRs (padj < 0.05). (E) A volcano plot showing the distribution of *Dnmt1-*KO DEGs, with significant genes (padj < 0.05) shown in dark blue, and DEGs with DMRs associated shown in purple. (F) *Egr1* DMR (qval < 0.05, gray vertical bar) in *Dnmt1*-KOs (purple) compared to control NPCs (orange). Flanking regions of DMR are ±5kb. On the x-axis, each tick mark represents a CpG in the mouse genome. (G) RNA-seq results showing normalized *Egr1* counts in *Dnmt1*-KOs compared to control NPCs. Adjusted pval < 0.05 (H) RNA-seq results showing normalized *Zim1* counts in *Dnmt1*-KOs compared to control NPCs. Adjusted pval < 0.05 (I) *Gsx1* DMR (qval < 0.05, gray vertical bar) found in *Kmt2a*-KO NPCs (orange) compared to control NPCs (purple). Flanking regions of DMR are ±5kb. On the x-axis, each tick mark represents a CpG in the mouse genome. (J) RNA-seq results showing normalized *Gsx1* counts in *Kmt2a*-KOs and controls. *p < 0.05, **p < 0.01, ns = not significant.

Given the extensive DNAm changes in *Dnmt1*-KO, we explored whether *Dnmt1*-KO DMRs show enrichment of TF binding motifs using an Analysis of Motif Enrichment (AME; MEME-suite, mouse HOCOMOCO v11 core motifs). This analysis identified 170 known TF binding motifs that were significantly enriched (Fisher’s exact test, adjusted pval < 0.05, **Figure 2D and Supplementary Table 4**), including a motif for MECP2, which has been shown to recruit DNMT1 to hemi or fully methylated CpGs (Kimura & Shiota, 2003). However, we also found other noteworthy TF motifs, some with CpG sites in their binding motifs, such as HIF1α, AHR and ARNT, and others lacking CpGs in their motifs, including SMAD3, ESR2, NKX2-5 and MXI1 (**Figure 2D, Supplementary Table 4**). Several hypoxia-related TF motifs (HIF1α, EPAS1 and ARNT) emerged. These are of particular interest given our recent observation of dysregulated hypoxia signaling in Kabuki syndrome, another MDEM, caused by mutations in *KMT2D* (Carosso et al., 2019). We next asked if the enrichment of TF motifs in the *Dnmt1*-KO associated DMRs could reflect known DNMT1 interaction partners. We compared these motifs with known DNMT1 interactors from BioGRID (Oughtred et al., 2021) and found that 13 of the 24 interactors represented in the HOCOMOCO database exhibited motif enrichment in *Dnmt1*-KO DMRs (**Supplementary Figure 2A**).

To investigate whether methylation changes in *Dnmt1*-KOs coincide with changes in transcription, we performed RNA-seq on *Dnmt1-*KO cells. We identified 3,454 differentially expressed genes (DEGs) compared with controls (**Supplementary Table 5**). We used HOMER to annotate the *Dnmt1*-KO DMRs and found that 2,713 of the DEGs (78.5%) are associated with 26,909 of the DMRs (19%; **Figure 2E, purple**). Among these were *Ppfibp1* (**Supplementary Figure 2B**), a gene encoding a factor involved in synaptic protein-protein interactions (PPIs)(Wei et al., 2011), the immediate early genes *Jun*, *Fos* and *Egr1* (**Figure 2F-G, Supplementary Figure 2C)** (Kim et al., 2018; Veyrac et al., 2014), as well as *Rora* and *Itga8* (**Supplementary Figure 2C-D**), previously identified DNMT1 target genes (Tao et al., 2022). We also found that mouse imprinted genes were significantly overrepresented in the *Dnmt1*-KO associated DEGs (Fisher’s exact test, OR = 1.83, p = 0.0118), including *Zim1* (**Figure 2H**)(Kim et al., 1999).

We next examined the impact of knocking out KMT2A, a histone methyltransferase which is expected to have a more indirect influence on DNAm compared to DNMT1. Nevertheless, pathogenic variants in *KMT2A* are the cause of Wiedemann-Steiner syndrome (OMIM, 605130) which has a distinct DNAm signature in peripheral blood (Foroutan et al., 2022). To investigate whether *Kmt2a* loss also perturbs DNAm in neuronal progenitors, we performed ONT sequencing on *Kmt2a*-KO NPCs in duplicates, which revealed only three genome-wide significant DMRs compared to controls. One of these DMRs was located in the 3’UTR of the *Gsx1* gene (meanDiff = 0.001121656, qval = 0.042, **Figure 2I**). We also performed RNA-seq after *Kmt2a* KO and found 2,105 DEGs compared to controls (**Supplementary Table 6**). Among the previously identified DMRs, only one is associated with a gene exhibiting significant changes in gene expression, *Gsx1* (**Figure 2J**). GSX1 is a TF expressed early in NPCs and is known to promote maturation in specific neuronal subtypes (Coltogirone et al., 2023; Pei et al., 2011). Notably, a prior study reported reduced H3K4me1 near this DMR in *Kmt2a* knockdown neurons derived from a mouse model (Kerimoglu et al., 2017) (**Supplementary Figure 2E**), supporting that *Gsx1* may be a direct target of KMT2A in mNPCs. We then queried GSX1 targets genes in our *Kmt2a*-KO DEGs. Using the TFBS database, which predicts TF-target gene interactions from DNA motifs and ENCODE DNase I footprints (Plaisier et al., 2016), we identified 133 GSX1 target genes (19.2% of 693 expressed targets) that overlapped with *Kmt2a*-KO DEGs. However, this overlap was not significantly enriched (Fisher’s exact test, OR = 1.18 and p = 0.055). To further explore whether *Gsx1* regulation might also be influenced by other EM genes, we examined the *Dnmt1*-KO. While we identified two intergenic DMRs near (∼30K and ∼16.5K upstream) the *Gsx1* locus in *Dnmt1*-KO (**Supplementary Figure 2F-G**), *Gsx1* did not exhibit corresponding transcriptional changes in *Dnmt1*-KO. Interestingly, its downstream targets were significantly enriched amongst *Dnmt1*-KO DEGs (Fisher’s exact test, OR = 1.16, p = 0.044). Finally, in contrast to the *Dnmt1*-KO DEGs, we found no enrichment of imprinting genes in *Kmt2a*-KO DEGs (Fisher’s exact test, OR = 1.23, p = 0.295). Thus, although imprinting is regulated through chromatin state, the effect is likely EM-gene specific.

### Convergence of gene expression changes after *Dnmt1* and *Kmt2a* disruption

Given the phenotypic overlap among multiple MDEMs, we next investigated the overlap in molecular consequences between *Kmt2a-*KOs and *Dnmt1-*KOs. When we compared the DEGs from both *Dnmt1-*KO and *Kmt2a-*KO cells, we found a substantial overlap of 1,543 DEGs (44.7% of *Dnmt1*-KO DEGs and 73.3% of *Kmt2a*-KO DEGs), including 679 upregulated and 861 downregulated genes (Fisher’s exact test, OR=11.6, p < 0.0001, **Figure 3A, Supplementary Table 7**). Moreover, gene expression changes were highly correlated between the two datasets (Spearman’s ρ = 0.94, p<0.0001, **Figure 3B**), and almost all shared DEGs were concordant in regard to the direction of change. In both datasets, the upregulated genes were enriched in neuronal differentiation and maturation GO terms (**Figure 3C**), while the downregulated genes were enriched in GO terms linked to cell cycle and chromosome organization (**Figure 3D**). We also performed a gene set enrichment analysis (GSEA) of both datasets, which revealed significant enrichment of downregulated genes in Hallmark terms such as MYC and E2F targets, and G2M checkpoint pathways (FDR < 0.05, **Figure 3E, Supplementary Figure 3A-C**), consistent with reduced proliferative activity. While *Myc* itself, a master regulator of cell-cycle progression (Bretones et al., 2015), was not differentially expressed in either dataset (**Supplementary Figure 3D-E**), many canonical MYC downstream effectors, including E2F genes, Cyclins, *Cdk1*, *Cdk2*, *Cdkn2b/p15*, and *Cdkn1a/p21*, followed the expected direction of change (**Figure 3F, Supplementary Figure 3F**)(Bretones et al., 2015). MYC regulators such as *Usp28* and *Skp2* were also downregulated (**Figure 3F, Supplementary Figure 3F**). Given that both KMT2A and DNMT1 have reported physical interactions with MYC on BioGRID (Kalkat et al., 2018), these patterns suggest that disruption of either factor may drive cells towards cell-cycle exit and premature differentiation.

**Figure 3:**
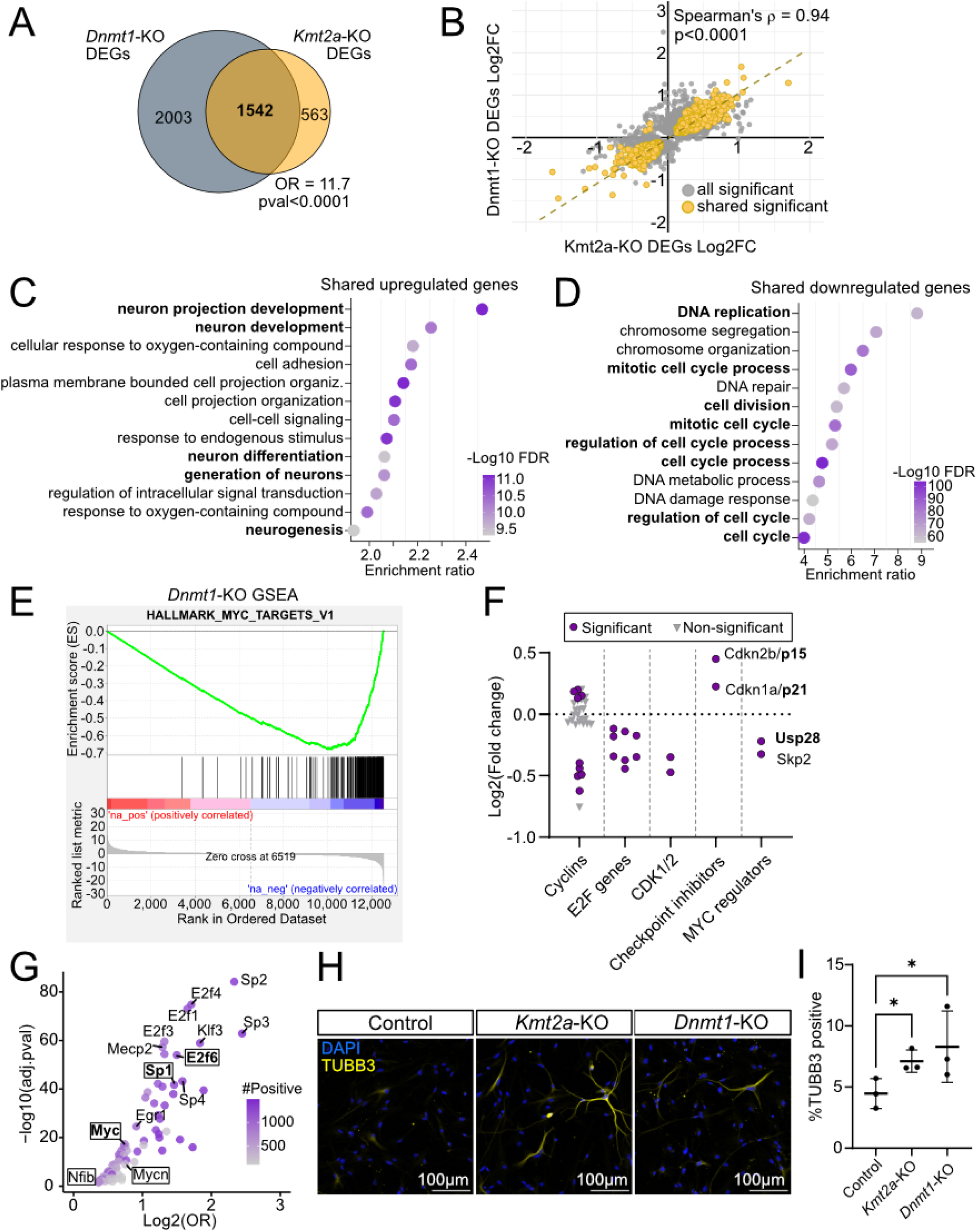
Premature differentiation in *Kmt2a*-KO and *Dnmt1*-KO. (A) A Venn diagram showing extensive overlap of DEGs in *Dnmt1*-KO and *Kmt2a*-KO cells compared to control NPCs, with results from Fisher’s exact test (OR = 11.7, p < 0.001). (B) Scatter plot showing Log2FC of significant DEGs from both *Dnmt1-*KOs and *Kmt2a-*KOs in gray, and the shared significant DEGs in yellow. Spearman’s correlation was calculated for the shared DEGs (R = 0.94, p-val < 0.0001). (C) GO term analysis (Biological Process) of overlapping upregulated genes, showing enrichment ratio of each GO-term on the x-axis and color depicting the FDR. (D) GO term analysis (Biological Process) of overlapping downregulated genes showing enrichment ratio for each GO-term on the x-axis and color depicting the FDR. (E) GSEA plots showing the MYC_targets_V1 Hallmark category for *Dnmt1*-KO genes. (F) RNA-seq results showing gene expression changes in *Dnmt1*-KO NPCs compared with controls. Y-axis shows Log2fold changes of genes belonging to the categories on the x-axis. Purple dots show significant genes; gray triangles show non-significant genes. (G) Scatter plot of TFs ranked by the number of promoters containing their binding motifs. The odds ratio indicates motif overrepresentation in promoters of shared DEGs. Boxes label shared BioGRID interactors. (H) IF staining of mNPCs after *Kmt2a-*KO and *Dnmt1*-KO compared with controls (blue = DAPI, yellow = TUBB3, magenta = EAAT2). (I) Quantification of the percentage of TUBB3 positive cells in *Kmt2a*-KO, *Dnmt1*-KO and controls. A one-way ANOVA with an FDR correction was used to calculate significance. *p < 0.05.

We also explored the expression of other EM factors and found that 27 EM genes were differentially expressed in both datasets (**Supplementary Figure 3G**), including *Ezh2*, which has previously been identified as a barrier to neuronal differentiation (Ciceri et al., 2024). Interestingly, both *Dnmt1* and a key interactor *Urhf1* (Liu et al., 2013) are significantly downregulated in both datasets. This suggests that core DNAm machinery is disrupted in both KOs although our data conclusively shows only a minimal effect on DNAm in the *Kmt2a-*KO and therefore establishes that the primary cause of the shared phenotypes is not due to changes in DNAm itself. Finally, *Trim28* and *Setdb1*, co-repressors of transposable elements (Fasching et al., 2015; Kato et al., 2018), were also found significantly downregulated in both datasets.

We next asked whether the shared DEGs were driven by common regulatory programs, using motif enrichment analysis of their promoter regions. Analysis of Motif Enrichment (AME; MEME-suite) of promoter regions of the shared DEGs identified common TF motifs, including members of the specificity protein (SP) and E2F families, most notably SP2 and E2F4 (**Figure 3G, Supplementary Table 8**), both of which are known regulators of cell cycle progression and differentiation in neuronal cells (Liang et al., 2013; Persengiev et al., 1999). We wondered if any of these TFs could interact with both KMT2A and DNMT1, thereby possibly explaining the overlapping expression changes. By comparing known interactors that were expressed in our system for either protein from BioGRID, we found a significant overlap of 53 interactors (Fisher’s exact, OR = 18.5, p<0.0001, **Supplementary Figure 3H**). Five of these had enriched motifs in the shared DEGs, most notably at SP1 and E2F6 targets, but also MYC (**Figure 3G, boxes**).

To determine whether *Kmt2a*-KO and *Dnmt1*-KO mNPCs undergo premature differentiation, we explored markers for neuronal and glial differentiation in both KOs and scrambled control mNPCs, using WB and IF staining. WB analysis revealed a significant increase in the neuronal marker TUBB3 in *Kmt2a*-KO mNPCs, but not the *Dnmt1*-KO mNPCs, compared to controls (**Supplementary Figure 4A**). No notable differences in the proliferation marker PCNA were detected in either KO model (**Supplementary Figure 4B**). Interestingly, IF staining demonstrated a greater number of TUBB3-positive cells in both *Kmt2a*-KO and *Dnmt1*-KO cultures, compared to scrambled controls, along with distinct morphological changes indicative of differentiation, such as smaller and elongated cell-bodies (**Figure 3H-I**). Increased expression of the astrocyte marker EAAT2 was also observed in *Dnmt1*-KO cultures, whereas no significant change was detected in *Kmt2a*-KO cells (**Supplementary Figure 4C-D**). Prior work has shown prematurely poised neuronal enhancers in primary mNPCs from a *Kmt2a*-haploinsufficient model, a phenotype which normalized after genetic rescue of *Kmt2a* (Reynisdottir et al., 2025). We found that NPCs from this model exhibit a similar premature neuronal differentiation phenotype as in our full KO (**Supplementary Figure 4E-F**), supporting that this is a feature of *Kmt2a* disruption. These findings support that loss of both factors contributes to premature differentiation of mNPCs but that they differ in impact. Specifically, loss of *Kmt2a* leads to a robust and uniform neuronal differentiation phenotype, whereas *Dnmt1* deficiency promotes both neuronal and glial differentiation, but potentially in a more variable manner.

### Evaluation of changes that occur during normal differentiation

Given the premature differentiation observed in EM-KOs, some of the expression changes may reflect differences in cell state composition across genotypes. To explore the extent of this in our EM-KOs, we differentiated wild-type (WT) mNPCs into mixed neuronal cultures (see methods) and demonstrated that these cultures express typical markers for neurons and astrocytes, as well as synaptic markers at Day 6 (**Supplementary Figure 5A**). We then performed RNA-seq on mNPCs, Day 2, and Day 6 differentiating cells (schematic in **Figure 4A**), which yielded 3706 and 3441 DEGs at Day 2 and Day 6, respectively, compared with mNPCs (**Supplementary Table 9-10**), and changes were consistent with developmental time-point (**Supplementary Figure 5B**). Both time-points were enriched in neurogenic and synaptic terms, consistent with ongoing neural differentiation and maturation of the culture (**Supplementary Figure 5C-D**). Additionally, a GSEA of cell-types showed significant enrichment in cell-type markers for the three main brain-cell lineages (neurons, astrocytes and oligodendrocytes, **Supplementary Figure 5E**), as would be expected.

**Figure 4:**
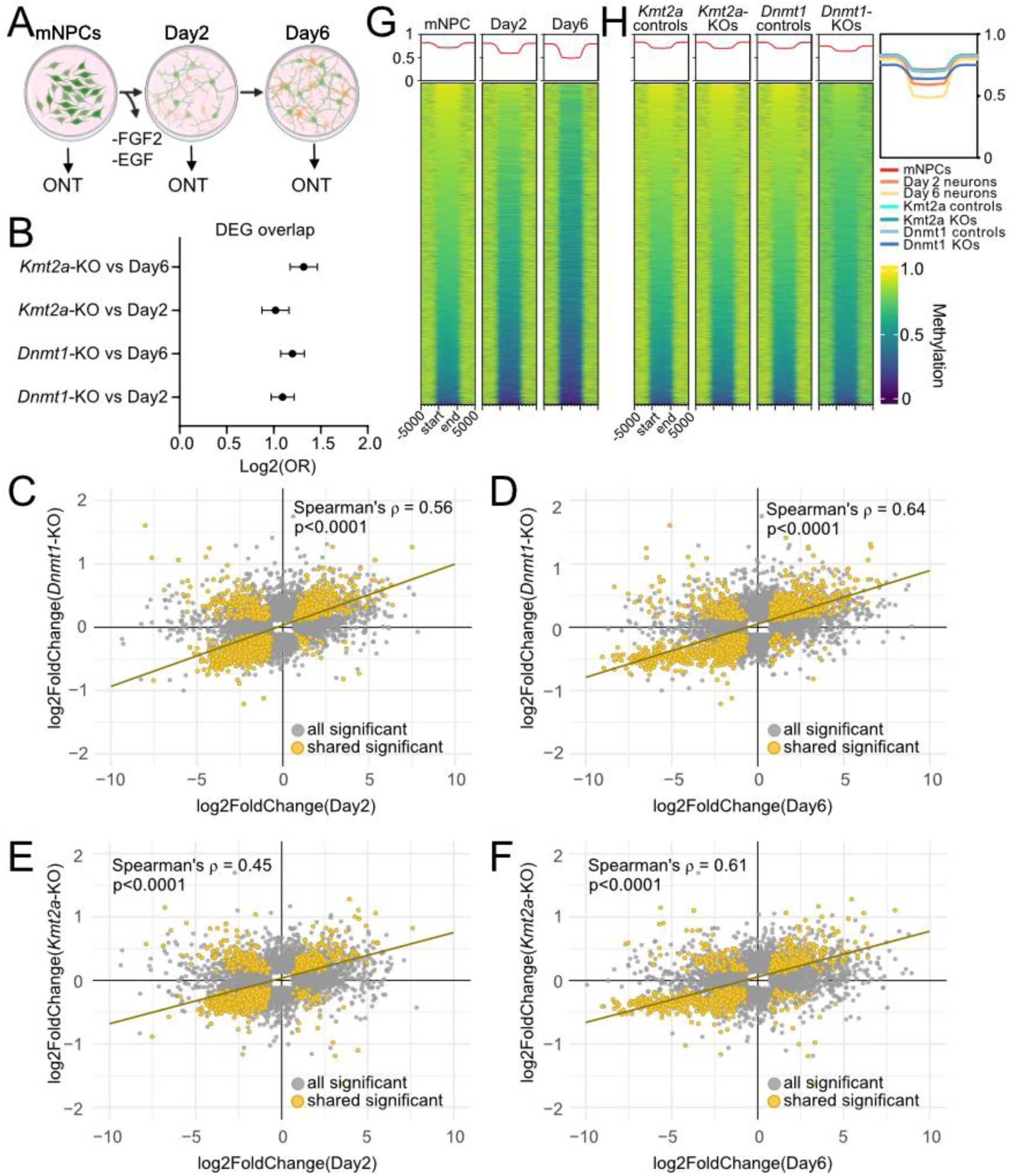
Changes observed in DNAm and gene expression during differentiation of NPCs. (A) A schematic of the neurodevelopmental model. (B) Odds ratio for the overlap between *Kmt2a*-KO and *Dnmt1*-KO DEGs with NPC-Day 2 and NPC-Day 6 DEGs, with 95% confidence intervals shown (Fisher’s exact test). (C-F) Significant Spearman correlations between DEGs of (C) *Dnmt1*-KO vs Day 2, (D) *Dnmt1*-KO vs Day 6, (E) *Kmt2a*-KO vs Day 2, (F) *Kmt2a*-KO vs Day 6. Axis represents log2(Fold change) for each dataset. (G) Day 6 DMRs are shown in mNPCs, Day 2 and Day 6 neurons. (H) Day 6 DMRs in KO and control samples. In the enriched heatmaps in (G) and (H), each line demonstrates the average smoothed methylation values across all CpGs in the corresponding DMR. DMRs were aligned at the DMR start site (start) and end site (end), ±5kb on either side. Yellow shows higher smooth methylation but blue shows lower smooth methylation values.

To understand the extent of transcriptional changes that may be secondary to premature differentiation in *Dnmt1-*KOs and *Kmt2a-*KOs, we compared DEGs from each KO to those identified during normal neuronal differentiation (mNPCs vs. Day 2 and Day 6). Interestingly, DEGs from both KOs showed strong and consistent overlap with normal differentiation associated gene expression changes (ORs > 2 in all comparisons, p < 0.0001, Fisher’s exact test; **Figure 4B**) and displayed a linear correlation (Spearman’s ρ > 0.45 for all comparisons, p < 0.0001; **Figure 4C-F**). These results suggest substantial convergence between KO-induced and differentiation-related transcriptional programs, consistent with a premature differentiation phenotype in the previously described EM-KOs.

To map the temporal landscape of DNAm during neuronal differentiation, we profiled genome-wide DNAm using ONT in mNPCs at Day 0, Day 2, and Day 6 of differentiation (schematic in **Figure 4A**). A principal component analysis (PCA) of genome-wide DNAm showed that the developmental stage was the main driver of methylation variance, as captured by PC1 (**Supplementary Figure 5F**). While differential methylation analysis identified no significant changes at Day 2 compared to Day 0 mNPCs, it revealed 8,627 genome-wide significant DMRs (qval < 0.05) at Day 6 compared to Day 0 mNPCs, indicating that most DNAm remodeling occurs relatively late in differentiation (**Figure 4G, Supplementary Table 11**). Genome-context analysis showed that on Day 6, similar to *Dnmt1*-KO DMRs, DMRs were enriched in introns, 3’ UTRs, and exons, but they were depleted in promoters, 5’ UTRs, and intergenic regions (**Supplementary Figure 5G**). To assess whether the methylation dynamics observed during normal differentiation are recapitulated in the KOs, we examined the methylation state of the Day 6 DMR regions in *Dnmt1*-KO and *Kmt2a*-KO cells. *Kmt2a*-KO cells maintained a methylation profile similar to undifferentiated mNPCs at these loci, while *Dnmt1*-KO cells exhibited a methylation pattern resembling that of Day 2 neurons (**Figure 4H**). However, methylation levels adjacent to these Day 6 DMRs were also lower in *Dnmt1*-KO cells, suggesting that the effects are not specific to those regions but rather a global effect. Additionally, 1,154 of the Day 6 DMRs were also identified in *Dnmt1*-KO cells (OR = 1.48, p < 2.2e-16, **Supplementary Table 12**). This suggests that the methylation changes over these Day 6 DMRs are driven by global hypomethylation in *Dnmt1*-KO cells but that they are also consistent with aspects of premature differentiation. Taken together, these results suggest that transcriptional changes precede widespread DNAm remodeling, supporting a model in which DNAm acts as a secondary layer that consolidates, rather than initiates, neurodevelopmental programs.

### An expanded cohort of EM-KOs shows subtle DNAm changes

We next broadened our analysis to assess DNAm abnormalities across a larger number of EM factors. Using ONT long-read whole genome sequencing, we profiled 44 additional EM-KO cells alongside matched scrambled controls. These data were pooled with previously generated data from the *Dnmt1-*KO and *Kmt2a-*KO, resulting in a total of 46 EM-KOs and 16 control samples. To systematically evaluate genome-wide DNAm changes, we partitioned the mouse genome into ∼2.7 million 1 kb tiles (each containing > 2 CpGs) and compared methylation levels across groups. A PCA plot of these tiles revealed that two known DNAm machinery components, *Dnmt1* and *Dnmt3b*, showed the most dramatic DNAm abnormalities (**Supplementary Figure 6A**). When these two were excluded from the analysis, *Kat6a*, *Chd5*, and *Tet1* emerged as outliers, clustering separately from the remaining EM-KOs (**Supplementary Figure 6B**). Despite this variation, no shared DMRs were identified across the 46 EM-KOs.

### Promoter DNAm alterations depend on interactions with the DNAm machinery

Given the diverse roles of EM genes in modulating DNAm, we hypothesized that shared DMRs may exist within specific subgroups of EM-KOs with related molecular functions. Therefore, classifying EM genes based on regional DNAm patterns could help uncover convergence. Promoter regions are well-established regulatory elements that are easily assigned to the corresponding genes and involved in transcriptional control (Deaton & Bird, 2011). To investigate whether a subset of EM-KOs exhibit similar DNAm changes at gene promoters, we applied non-negative matrix factorization (NMF) to cluster the 46 EM-KOs based on DNAm patterns at the promoters of 20,482 mouse protein-coding genes (**Supplementary Figure 6C)**. Hierarchical clustering of EM-KOs using the NMF-derived coefficients of latent factors further characterized three latent factors (**Figure 5A**). Specifically, cluster 1 included *Chd3, Setd4, Setd1a, Setd1b, Kdm5b*, and *Kdm1a*, showing the highest coefficient in latent factor 3. In contrast, cluster 2 included the controls and showed low coefficients in all three latent factors. Finally, cluster 3 included the genes *Kmt2a*, *Dnmt1*, *Chd1*, *Kdm6a*, *Hdac6*, *Ezh2*, and *Crebbp* and showed the highest coefficient in latent factor 2 (**Figure 5A**). *Dnmt3b-*KO showed the highest coefficient in latent factor 1 and did not cluster with other EM genes (**Figure 5A**). Interestingly, genes with the highest weight in latent factor 2 were significantly enriched in GO terms related to generation of neurons and neurodifferentiation (**Figure 5B**) as well as the Wnt signaling pathway (**Supplementary Figure 6D**). Conversely, no significant enrichment in GO terms was found for the other latent factor groups. However, separate DNAm analysis comparing each of these clusters to controls did not reveal any genome-wide significant DMRs. In the two clusters that did not group with controls (clusters 1 and 3), nearly all the gene members (excluding SETD4) had STRING defined interactions with the maintenance methyltransferase DNMT1 (**Figure 5A**). Indeed, components of these clusters had a significant enrichment in interactions with all the enzymatic DNAm machinery components (**Figure 5A, C**). This enrichment was strongest for the maintenance methyltransferase (DNMT1) but less so with *de novo* methyltransferases (**Figure 5C**). Furthermore, the same groups (Clusters 1&2 and DNMT3B), had a significantly higher number of STRING interactions with other members of the DNAm machinery (n = 25) (Boukas et al., 2019) compared to what was seen in cluster 2 (**Figure 5D**, one-sided Wilcoxon rank-sum test, W = 349.5, p-value = 0.0014). Additionally, these same groups all involve factors with enzymatic activity, while readers of epigenetic marks are only present in cluster 2. These results suggest that enzymatic EM factors, outside the core DNAm machinery, may influence promoter DNAm through interactions with components of the DNAm machinery.

**Figure 5:**
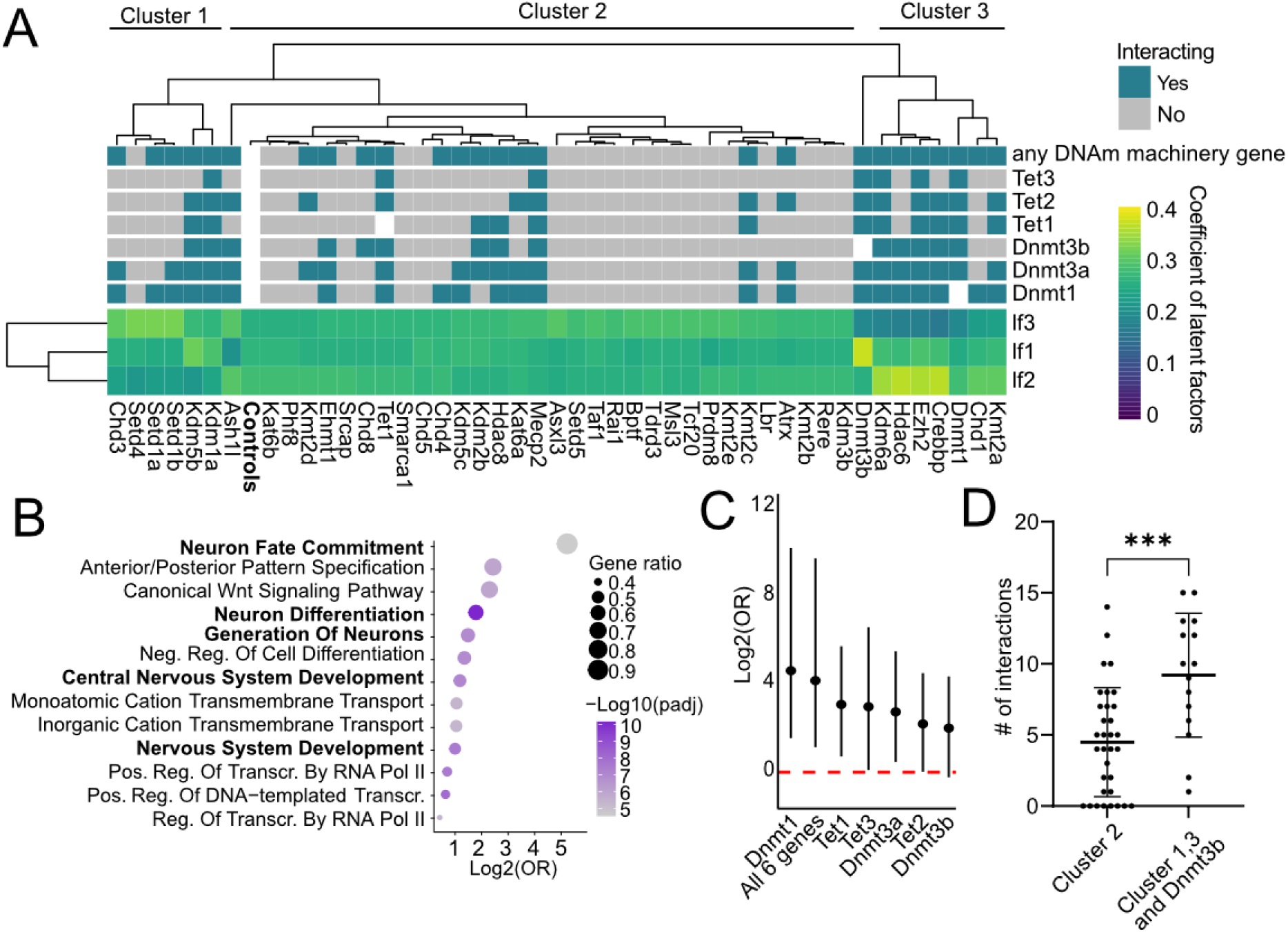
DNAm analysis of 46 EM-KO genes in NPCs reveals shared promoter effects. (A) A composite graph showing STRING interactions of EM components with enzymatic DNAm components on top, the heatmap showing coefficient of latent factors for each EM-KO across three latent factors. The color scale represents the coefficient of each latent factor (yellow shows higher but blue a lower latent factor coefficient). (B) Gene ontology (GO) term enrichment (Biological Process) of the genes associated with latent factor 2. Size represents the ratio of genes in the relevant GO term, color indicates the -log10(adjusted p value). (C) Forest plot showing enrichment of enzymatic DNAm machinery-interacting EMs in two relevant clusters compared to rest. Significance testing was done with a Fisher’s exact test. Dots represent Fisher’s exact test OR (log₂ scale), with lines indicating 95% confidence intervals. The horizontal dotted red line represents statistical significance. (D) Comparison of number of STRING interactions with 25 DNAm machinery components between the clusters. A Wilcoxon rank sum test was used for significance. lf1 = latent factor 1, lf2 = latent factor 2, lf3 = latent factor 3. ***p < 0.001, ns = not significant.

### Haplotype-specific DNAm within the context of EM-KOs

We used mNPCs derived from F1 offspring of the two strains (FVB maternal × B6J paternal, **Figure 1A**). As the B6J and FVB mouse strains differ genetically, with a SNV between the two genomes roughly every 630bp, it is possible to phase sequenced reads and perform an allele-specific methylation analysis. This is an average statistic, in practice roughly half of the two genomes can be phased. PCA of the phased data shows the same pattern between KOs and controls as for the unphased data (**Supplementary Figure 7A**) but supports the presence of haplotype-specific methylation between the two strains. Indeed, within the control samples, we identified 1,074 haplotype-specific DMRs (FWER < 0.05) (**Figure 6A, Supplementary Table 13**) (Do et al., 2016; Tycko, 2010), including one located in the exon 4 of *Tet3* (**Figure 6B**). Due to the consistent parental origin of the two genomes in these F1 crosses, imprinted loci will manifest as allele-specific methylation patterns within a single cross direction. Using 23 known ICR regions (Gigante et al., 2019), we have coverage on both haplotypes in 16 of the 23, and all 16 exhibit strong methylation differences between the two alleles (example in **Supplementary Figure 7B**).

**Figure 6:**
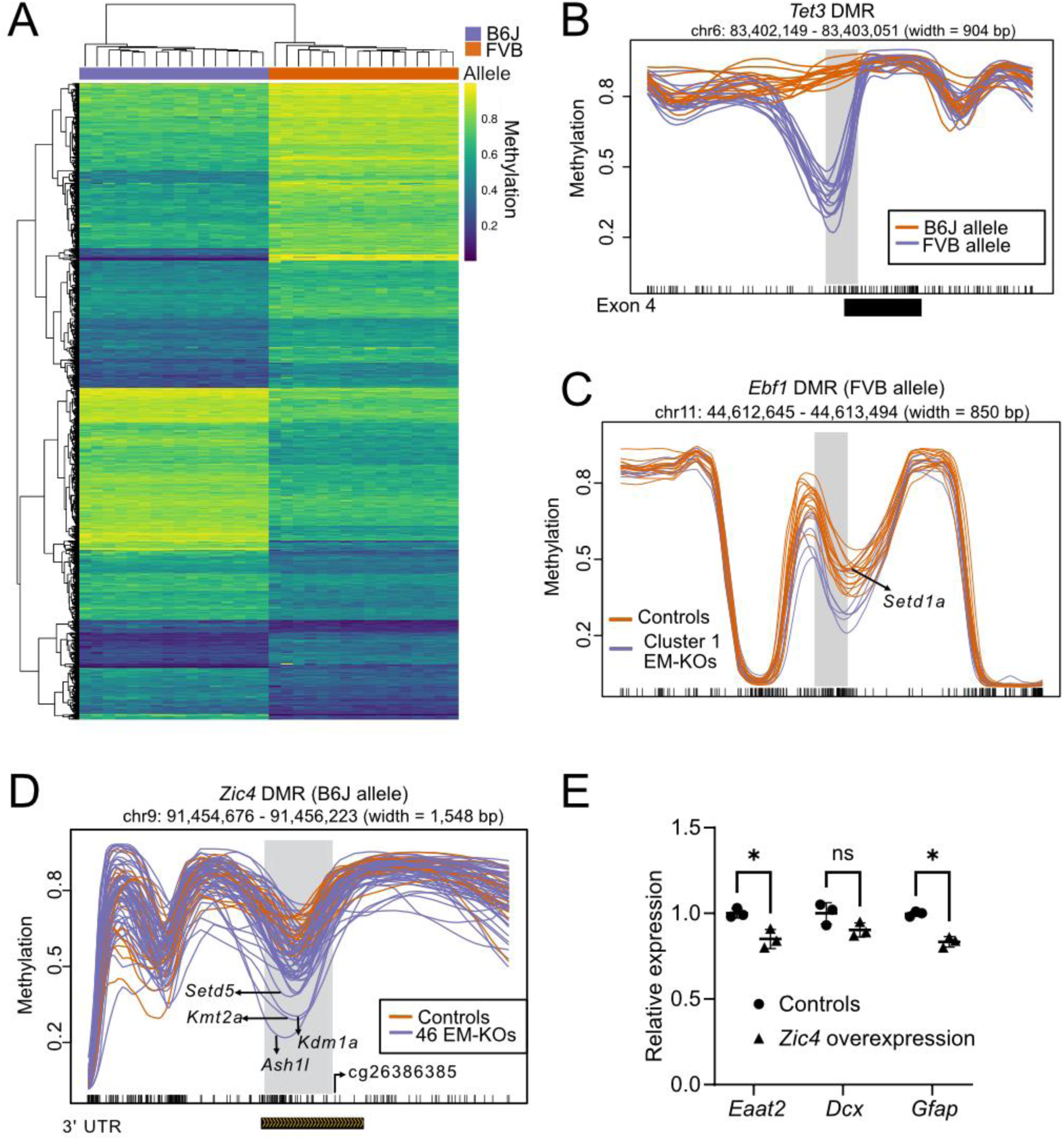
Haplotype-specific changes of DNAm. (A) A heatmap plot showing smoothed methylation of 1074 haplotype-specific DMRs across B6J and FVB haplotypes. Yellow and blue represent higher and lower DNAm, respectively. Each line in y-axis represents a single DMR across haplotyped controls but x-axis represents haplotyped controls. (B) A haplotype-specific DMR in the *Tet3* gene. (C) A line plot showing a DMR in 5’ UTR of *Ebf1* between the FVB allele of control mNPC (orange) and the FVB allele of Cluster 1 EM-KOs (purple). The DMR is highlighted in gray. Flanking regions of DMR are ±5kb. On the x-axis, each tick mark represents a CpG in the mouse genome. (D) A line plot showing a DMR in 3’ UTR of *Zic4* between the B6J allele of control mNPC (orange) and the B6J allele of 46 EM-KOs (purple). The DMR is highlighted in gray. Flanking regions of DMR are ±5kb. On the x-axis, each tick mark represents a CpG in the mouse genome. (E) WB analysis of Myc-ZIC4 protein after overexpression. (H) qPCR results showing relative mRNA levels of *Eaat2, Dcx* and *Gfap* after ZIC4 overexpression, compared with Myc-only controls; significance calculated using a t-test.

To determine if the EM-clusters, identified via the NMF analysis (**Figure 5A**) yielded any shared haplotype-specific DMRs, we performed a grouped analysis on the phased data for each of these clusters, compared to the controls. Cluster 1 (*Chd3*, *Setd4*, *Setd1a*, *Setd1b*, *Kdm5b*, and *Kdm1a*) showed a shared hypomethylated DMR localized to the FVB allele 4.6 kb upstream of the gene *Ebf1* (FWER = 0.03, **Figure 6C**), which encodes a TF involved in early neurogenesis (Garel et al., 1999; La Manno et al., 2021). In contrast, cluster 2 and 3 did not yield any shared DMRs compared to controls. Finally, we searched for haplotype-specific DMRs associated with the entire group of 46 EM-KOs and identified a single shared hypomethylated DMR localized to the B6J allele (FWER<0.05). This DMR is located in the 3’ UTR of *Zic4* (**Figure 6D, Supplementary Figure 7C**), a TF that plays a role in cerebellar development (Grinberg et al., 2004). Studies using DNA methylation arrays in humans, have shown that (a) the 3’ UTR of this gene is dynamically methylated during differentiation from hESC to neurons (Samara et al., 2022) and (b) methylation of the gene body contributes to an episignature for Sotos syndrome (Choufani et al., 2015). Although *Zic4* was not differentially expressed in *Dnmt1*-KO nor *Kmt2a*-KO mNPCs compared to scrambled controls (**Supplementary Figure 7D-E**), it was significantly downregulated (log2FC = -1.978, adjusted p-val = 0.00197) at Day 6 compared to Day 0 mNPCs in our neurodevelopmental model (**Supplementary Figure 7F**). To evaluate the functional role of ZIC4 in our system, we overexpressed Myc-tagged *Zic4* in mNPCs and compared with the Myc-tag alone. The results showed reduced expression of neuronal maturation markers (**Figure 6E, Supplementary Figure 7G**), suggesting that ZIC4 may maintain the NPC state and raising the possibility that it may act as a brake on neuronal maturation.

## Discussion

Our study presents a comprehensive investigation into how EM disruption affects DNAm and gene expression in a disease-relevant neuronal system, offering insights into the presence and functional impact of DNAm changes beyond peripheral blood. By combining a mNPC model with ONT sequencing, haplotype phasing, and transcriptome profiling, we characterized the functional consequences of 46 EM gene KOs. Disruption of *Dnmt1* leads, as expected, to extensive changes in DNAm. A PCA of genome-wide DNAm suggests that the second largest effect on DNAm is from *Dnmt3b* and, interestingly, *Kat6a*, although further investigation is required to assess these changes genome-wide. Since we aimed to identify DNAm changes that were likely to be causative rather than a consequence of the disease phenotype, we isolated and profiled the methylation levels relatively soon after KO (4 days). However, across the 46 EM-KOs, we observe little-to-no shared changes in DNAm and little evidence for KO-specific changes with the exception of a few KOs such as *Dnmt1* and *Dnmt3b*. This does not exclude the possibility that DNAm changes could occur at a later time point. Our data suggests that multiple TFs such as GSX1, MYC, SP and the E2F family TFs may mediate or interpret EM-related epigenetic disruption. For instance, *Gsx1* was differentially expressed in *Kmt2a*-deficient cells, and many of its target genes were dysregulated in both *Kmt2a*-KOs and *Dnmt1*-KOs. Similarly, MYC, SP and E2F family TF binding motifs were enriched in promoters of shared DEGs, and GSEA showed MYC targets significantly negatively enriched. Interestingly, MYC, SP1 and E2F6 specifically are known binding partners of both KMT2A and DNMT1. These findings point to a model in which the EM indirectly modulates transcriptional networks through TFs, either by altering DNAm at TF binding sites or via interactions with the TFs themselves. Similarly, our clustering of promoter DNAm suggests that, at least in some EM contexts, physical interactions with the DNAm machinery plays a role. Importantly, some TFs such as EGR1 are known to recruit TET1, further linking transcriptional networks to active demethylation pathways (Sun et al., 2019).

Interestingly, we identified a hypomethylated DMR in the 3’ UTR of *Zic4* which was shared across all 46 EM-KOs in one of the two haplotypes. This gene is implicated in both murine and human neurodevelopment and DNAm changes in the gene has been identified in peripheral blood from individuals with Sotos syndrome, a neurodevelopmental disorder caused by mutations in the epigenetic regulator *NSD1* (Choufani et al., 2015). However, the methylation changes are modest, and this gene is not differentially expressed in our EM-KOs. Nevertheless, *Zic4* is downregulated during differentiation and overexpression of *Zic4* reduced expression of markers for neuronal and glial differentiation (**Supplementary Figure 7F, Figure6E**) suggesting that it may play a role in maintaining the NPC state. It is not clear whether this gene is important in explaining the phenotypic overlap in the MDEM group, but it could nevertheless have potential utility as a biomarker.

We have established a premature differentiation phenotype arising because of either *Dnmt1-*KO or *Kmt2a-*KO. This is something we and others have observed as a consequence of disrupting several other EM factors. For example, in KMT2A-deficient mice we previously found H3K4me1 enrichment at neuronal enhancers in undifferentiated NPCs (Reynisdottir et al., 2025), suggesting that these enhancers show premature poising. Additionally, prior work on Kabuki syndrome, another MDEM caused by mutations in *KMT2D*, the same phenotype has been reported in two disease relevant contexts (Carosso et al., 2019; Fahrner et al., 2019). Moreover, studies of Kabuki syndrome neurons from a mouse model have shown an overrepresentation of aging-related loci (Boukas et al., 2024), further suggesting that premature differentiation may represent a unifying phenotype across a subset of MDEMs. Interestingly, in our NMF analysis, KMT2A and DNMT1 clustered with EZH2, KDM6A and CREBBP, all of which have been shown to regulate neuronal stem cell maintenance and differentiation (Lei & Jiao, 2018; Pereira et al., 2010; Wang et al., 2010). Genes associated with the latent factor of this group were enriched for functions relating to neuronal differentiation (**Figure 5B**). This could suggest that even though the DNAm changes are very small, the disruption of these EM factors affects a core NPC gene network.

Despite the modest DNAm changes, we observed an impressive convergence of transcriptional dysregulation between *Dnmt1*- and *Kmt2a*-KOs, along with the premature differentiation phenotype. Comparing the transcriptional changes in these two EM-KOs to transcriptional changes during normal neuronal differentiation, we conclude that part of the overlap is secondary to the premature differentiation. Nonetheless, we note that the overlap between *Dnmt1-*KO and *Kmt2a-*KO cells appears stronger (larger OR) than either of the overlaps with normal differentiation. Combined with the lack of methylation changes in *Kmt2a-*KOs, we have shown that the expression changes are not a consequence of DNA methylation changes and are more likely to drive this particular phenotype.

Our data also points to a coordinated disruption of MYC-dependent regulatory programs after *Kmt2a*- and *Dnmt1-*KO. Although *Myc* transcript levels were unchanged, both KMT2A and DNMT1 are known MYC interactors (Kalkat et al., 2018), and we observed consistent downregulation of MYC target genes in the expected direction (**Figure 3E-F**). This pattern strongly suggests reduced MYC activity. As MYC is a master cell-cycle regulator (Dong et al., 2014; García-Gutiérrez et al., 2019), either the loss of interaction with DNMT1 or KMT2A or the downregulation of its deubiquitinator USP28 (Popov et al., 2007), might be a critical factor contributing to the premature cell-cycle exit and neuronal differentiation phenotypes we observe. Further work is required to understand how the shared expression changes arise, and especially to identify which epigenetic changes, if any, are causal in this context. Furthermore, differentiation state should be carefully considered when analyzing gene expression, since premature or altered differentiation can be a major source of the observed changes and thus a confounder of such studies.

Beyond providing mechanistic insight, this work establishes a framework for using long-read sequencing, haplotype phasing, and transcriptomic profiling in disease-relevant cells to understand the role of epigenetic regulators in health and disease.

## Materials and methods

### Animals

B6J.129(Cg)-Gt(ROSA)26Sortm1.1(CAG-cas9*,-EGFP)Fezh/J(Strain#026179) (B6J) and FVB.129(B6)-Gt(ROSA)26Sortm1.1(CAG-cas9*,-EGFP)Fezh/J(Strain#026558) (FVB) mice, each carrying a heterozygous Cas9 transgene, were obtained from Jackson laboratory. Mice were bred within their respective background strains (B6J and FVB) to generate homozygous Cas9-expressing offspring. DNA was isolated from ear clips, and Polymerase Chain Reaction (PCR) was used to confirm the Cas9 cassette using primers design provided by the Jackson laboratory (**Supplementary Table 14**). All mice were kept on a 12 h dark-light cycle at room temperature (21-23°C) with relative humidity around 40% and maintained with ad libitum diet and water. Mouse importation and experimental protocols were in accordance with Icelandic Food and Veterinary Authority (license no. 2208602) and approved by the National Expert Advisory Board on Animal Welfare.

### Dissections and primary NPCs isolation

The NPCs were isolated from an F1 hybrids of homozygous Cas9 mice (B6J and FVB). The isolation procedure was based on published protocols (Bernas et al., 2017; Rybachuk et al., 2019) with minor modifications, see Supplementary Materials for details. NPCs were grown in neuronal growth medium (NGM), which included Neurobasal medium supplemented with 1% B27 supplement (ThermoFisher, 17504044), 1% Glutamax (ThermoFisher, 35050061), 0.25% Penicillin-streptomycin (ThermoFisher, 15140122) 20 ng/ml EGF (PeproTech, AF-100-15), 20 ng/ml FGF2 (PeproTech, 100-18B) and 2 µg/ml Heparin (MP Biomedicals, 0210193125) on PDL/Laminin coated 6-well plates, and frozen down in NGM + 10% DMSO (Santa Cruz, sc-358801).

### Neuronal progenitor differentiation

NPCs (passage 3-4) were differentiated into mixed neuronal cultures by collecting them as previously described, counting and seeding 150.000 cells/well on a PDL/Laminin coated 24-well plate in 0.5 ml N2-Neuronal Induction Medium (N2NIM) consisting of Neurobasal Plus Medium (ThermoFisher, A3582901) with 1x B-27 Plus Supplement (ThermoFisher, A3582801), 1x CultureOne Supplement (ThermoFisher, A3320201), 1x N2 Supplement (ThermoFisher, 17502048), 0.25x Penicillin-streptomycin and 0.25x GlutaMax. This was supplemented with 20ng/ml FGF2 and 10ng/ml BDNF (Peprotech, 450-02). On day 2 and 4, the medium was changed to N2NIM supplemented with 10ng/ml FGF2, 10ng/ml BDNF and 10ng/ml GDNF (ORF Genetics, IK0900). At Day2 and Day6 of differentiation, neuronal cultures were collected and DNA and RNA isolated as described below.

### Guide RNA design and isolation of plasmids

CRISPR/Cas9 guide RNAs (sgRNAs) were designed with an online tool (https://chopchop.cbu.uib.no/) (Labun et al., 2019). Two sgRNAs, with the highest predicted efficiency and minimal mismatches per gene, were selected to ensure out-of-frame upon Cas9-mediated cleavage. All sgRNAs were verified with BLAT to ensure that none target any other sequences in either of the two genomes. Eight scrambled control sgRNA pairs (random sequence not binding in the murine genome) were designed as controls. Lentiviral CRISPR vectors (Replogle et al., 2020), each including one pair of sgRNAs (**Supplementary Table 1**) were synthesized by Vectorbuilder. Bacterial stabs were streaked onto LB plates and plasmids isolated using ZymoPURE II plasmid Midiprep kit (Zymo, D4201) and stored at -20°C until needed. All 54 dual guide-RNA lentiviral plasmids were Sanger sequenced to confirm the presence of the correct sgRNA pairs.

### Cell culture

For virus production, HEK293T cells were cultured in culturing media, DMEM/F12 supplemented with GlutaMAX™ (ThermoFisher, 31331093) and 10% Fetal Bovine Serum (FBS) (ThermoFisher, 10270106). All cells were incubated at 37°C with 5% CO_2_. HEK293T cells were frozen down in culture media supplemented with 10% DMSO.

### Virus production and transduction into NPCs

HEK293T cells in a T75 flask were transfected to produce lentivirus. The transfection mixture contained 2.29 µg lentiviral CRISPR plasmid, 2.08 µg psPAX2 (Addgene, 12260), 0.63 µg pMD2.G (Addgene, 12259), combined with 15 µl Lipofectamine 3000 and 10 µl P3000 (ThermoFisher, L3000015), and brought to a final volume of 250 µl with Opti-MEM™ (ThermoFisher, 31985047). The transfected HEK293T cells were cultured for 72 hours, and viral particles collected from the medium after 48 and 72 hours, passing the supernatant through a 0.45 µm syringe filter (Santa Cruz, sc-358814) to remove cell debris. After the second collection, 4× Lentivirus concentrator solution was prepared following the protocol from the MD Anderson Cancer Center Functional Genomics Core, and was used to precipitate viral particles (Kutner et al., 2009; Lo & Yee, 2007). Fifty µl viral aliquots were stored in - 80℃ until use. Transduction efficiency for each virus was tested in NPCs at several concentrations, using serial dilutions of the virus together with 8 µg/ml polybrene (Santa Cruz, sc-134220). Transduction efficiency was determined using fluorescence-activated cell sorting (FACS) of BFP fluorescent cells in NPC cells, 48 hours after transduction, and an efficiency of ∼30% was used to conduct all experiments.

### EM-KO in NPCs and validation

For EM-KO experiments, the NPCs were seeded on a 6-well plate 24 h prior to transduction. Cells were transduced with lentivirus in the presence of 8 µg/ml polybrene. Each lentivirus targeted a single EM gene and non-targeting controls were used as negative controls for viral infection and culturing effects. After 24 h, cells were split and selected by 0.5 µg/ml puromycin (ThermoFisher, A1113802) for 2 days. Cells surviving puromycin treatment were cultured for another day without puromycin to allow them to recover, then harvested for experiments using Accutase as described above. Timing was selected based on the fact that average protein half-life in mouse primary neurons is ∼86 hours (Mathieson et al., 2018) and for EM proteins it is considered even less (∼40 hours), thus most of the EM proteins would be expected to be degraded at the time of our evaluation. The KO efficiency was determined by PCR, WB and IF staining.

### RNA isolation, cDNA synthesis and Real Time Quantification PCR (RT-qPCR)

Cultured cells were harvested and lysed in TRIzol™ Reagent (ThermoFisher, 15596018). Total RNA was isolated using Direct-zol™ RNA Microprep Kit (Zymo Research, R2062) based on the manufacturer’s protocol. The concentration and quality of RNA was evaluated with Nanodrop (ThermoFisher). One µg total RNA was used for cDNA synthesis with High-Capacity cDNA Reverse Transcription kit (ThermoFisher, 4368814). RT-qPCR reactions were performed using Luna® Universal qPCR Master Mix (NEB, M3003) and a CFX384 Touch Real-Time PCR Detection System (BIO-RAD), according to manufacturer’s protocol. Any primers used are provided in **Supplementary Table 14.**

### Protein isolation and WB

NPCs were collected using Accutase and counted with a Countess II FL Automated Cell Counter (ThermoFisher). Cells were lysed at 15000-20000 cells/µl in 1x sample buffer (60mM Tris HCl pH 6.8, 2% SDS, 10% Glycerol, 0.01% Bromophenol blue, 1.25% Beta-me) supplemented with 1:100 Halt™ Protease Inhibitor Cocktail (ThermoFisher, 78437), 1:100 phosphatase inhibitor Cocktail (Cell signaling, 5870) and 1:100 PMSF Protease Inhibitor (ThermoFisher, 36978). The lysate was heated at 95°C for 5 min and 1:100 Benzonase nuclease (Sigma-Aldrich, sc-202391) added to degrade genomic DNA. The lysate was aliquoted and stored in -80°C. Western blotting was performed as has previously been described (Mahmood & Yang, 2012), where 15-25 µg of protein was loaded per sample, blotted onto PVDF membrane (ThermoFisher, 88520) and stained overnight using the following primary antibodies: rabbit anti-CHD1, 1:1000 (Cell signaling, 4351), mouse anti-ACTIN, 1:1000 (Abcam, ab8224), rabbit anti-EAAT2, 1:1200 (Abcam, ab41621), mouse anti-beta III Tubulin, 1:800 (Abcam, ab78078), rabbit anti-DNMT1, 1:1000 (Proteintech, 24206-1-AP), mouse anti-VINCULIN, 1:1000 (Sigma, V9131), rabbit anti-Myc-Tag, 1:1000 (Cell signaling, 2272). Mouse anti-PCNA, 1:7000 (Proteintech, 60097-1-Ig) was incubated at RT for 1.5 h. Blots were stained with secondary antibodies: Donkey anti-Mouse IgG, 1:15000 (LICOR, 926-68072), Donkey anti-Rabbit IgG, 1:15000 (LICOR, 926-32213) for 1h at RT. After washing, blots were visualized with Odyssey® CLx Imager and protein quantification was evaluated by Image J (Schindelin et al., 2012).

### Immunofluorescence

NPCs were cultured on PDL/Laminin-coated 8-well culture slide (Falcon, 354118) for 24-48 hours, fixed with 4% paraformaldehyde (PFA), blocked with 5% NGS in 0.1% PBS-T and stained using primary antibodies against rabbit anti-KMT2A, 1:200 (Cell signaling, 14197), rabbit anti-EAAT2 1:400 (Abcam, ab41621) and mouse beta III Tubulin 1:200 (Abcam, ab78078), overnight. The following day, cells were stained using secondary antibodies: Alexa goat anti-rabbit 647, 1:1000 (ThermoFisher, A-21244) and Alexa goat anti-mouse 555, 1:1000 (ThermoFisher, A-32727). Cells were mounted with Fluoromount-G™ Mounting Medium, with DAPI (ThermoFisher, 00-4959-52). Images were captured with Olympus Confocal microscopy (Olympus Fluoroview, FV4000), processed with Fiji and quantified with CellProfiler. Statistical analysis was done using GraphPad Prism v10.4.

### Overexpression

NPCs were seeded at a density of 250,000-300,000 cells per well in PDL and Laminin coated 6-well plates, 24 hours prior to transfection. The following day, cells were co-transfected with 2.5 µg of either the Myc-tagged *Zic4* overexpression plasmid (Origene, MR216949) or the corresponding Myc-alone plasmid backbone (control), along with 7.5 µl of Lipofectamine 3000, following the manufacturer’s protocol. Cells were harvested 24 hours post-transfection for downstream analysis.

### DNA sample isolation and preparation

Genomic DNA isolated from cells using Monarch® HMW DNA Extraction Kit for Cells & Blood (NEB, T3050) followed the manufacturers protocol for DNA extraction from cells. The concentration and quality of DNA was measured by Nanodrop (ThermoFisher) and Qubit to quantify the precise amount of double stranded DNA via Qubit dsDNA HS assay kit (ThermoFisher, Q33231). High molecular weight (HMW) DNA was sheared to 20-30 kb to achieve consistent size and to optimize ONT sequencing yield. HMW DNA was loaded to a hydro-tube assembled with hydropore-syringe (Diagenode, E07010003), inserted into the cassette, installed on a base unit and then sheared by Megaruptor® 3 (Diagenode) using the following settings: a volume of 70 µl, a speed of 30, and a concentration of 50 ng/µl. The size of sheared DNA was evaluated with HS Large Fragment 50 kb kit (Agilent, DNF-464-0500) on a Fragment Analyzer, proceeded by High Sensitivity Large Fragment 50 kb Analysis software.

### ONT-DNA sequencing library construction

Library preparation for ONT-DNA sequencing used the ONT Ligation Sequencing kit V14 (ONT, SQK-LSK114), using 1 µg sheared genomic HMW DNA in 47 µl nuclease-free water (ThermoFisher, Am9937) as input, followed by library preparing and loading R10.4.1 flow cell (FLO-PRO114M, ONT) on a PromethION 24 sequencer following the manufacturer’s instructions, see supplementary methods for details. Nanopore sequencing was performed using the following settings: a Run limit of 96 h, a Minimum read length of 200 bp, and Super-Accurate Basecalling Model with 5mC and 5hmC modified base context. To increase the output, flow cells were washed once after 22-24 h of sequencing using Flow Cell Wash kit XL (ONT, EXP-WAS004-XL), followed by 1-1.5 h waiting time. Then priming and loading library procedures were the same as described previously. Around 70 Gigabases estimated bases with an average read length of 20 kb were achieved per sample.

### Detection of KO efficiency from ONT sequencing data

We developed a computational tool, *samCRISPR*, to estimate CRISPR/Cas9 KO efficiency using ONT long-read whole-genome sequencing (WGS) data. The name “samCRISPR” references both SAMtools, from which the mpileup format is used in KO efficiency calculations, and CRISPR-Cas9 genome editing technology. SAMtools is used to sort, index, and filter aligned reads, and to generate mpileup output at the target site, which provides base-level resolution of read alignments. This output is parsed by our tool, *samCRISPR*, to identify substitutions, insertions, and deletions based on mpileup letter codes, allowing precise quantification of editing events and KO efficiency. The tool identifies sequencing reads that carry CRISPR-induced events within designated target regions using the mpileup output from SAMtools (Danecek et al., 2021) (**Figure 1F**). To eliminate the chance of counting sequencing errors in the vicinity of target regions as actual CRISPR events, the counting of CRISPR events was restricted to 1bp upstream and 1bp downstream of Cas9 cut sites. CRISPR events were defined as mutations occurring within ±1 base pairs of the predicted cut site, located 3 bp upstream of the PAM sequence. This narrow window was chosen to minimize false positives caused by sequencing or alignment errors near Cas9 cleavage sites. KO efficiency was calculated as the ratio of reads with CRISPR events in the target region to the total number of reads mapped to that region. To validate samCRISPR results, KO efficiencies for selected EM-KOs were manually inspected using the Integrative Genomics Viewer (IGV) (Robinson et al., 2011). *samCRISPR* accepts a BAM file, a reference FASTA, or BED-formatted sgRNA coordinates as input, and supports batch analysis of multiple BAM files and sgRNAs. It requires a Linux environment with SAMtools for mpileup generation and R computing environment for calculating Wilson score confidence intervals (Agresti & Coull, 1998).

### Phasing of ONT data and obtaining CpG-level methylation data

We utilized a custom graph genome-based approach, termed PLASMA (Pipeline for Long-read Allele-Specific Methylation Analysis), to integrate both FVB and B6 genomic sequences into a unified non-linear reference coordinate suitable for allele-specific methylation analysis with minimal introduction of reference bias (Davidovich, et al., in preparation). A genome graph, a structure composed of nodes and edges, captures alternative alleles, structural variants, and indels across multiple genomes (Liao et al., 2023; Zhang et al., 2025). Unlike linear references, genome graphs support the simultaneous representation of multiple haplotypes, making them particularly suited for admixed or hybrid genomes.

The whole-genome alignment between the FVB and B6J reference genomes was performed using MUMmer4 v4.0 (Marcais et al., 2018) to identify heterozygous variants. Next, Jellyfish (Marcais & Kingsford, 2011) was used to enumerate all 45-mers in each genome, which were subsequently classified into conserved nodes and variant edges. Edges shorter than 3 kb were aligned between genomes to derive CIGAR strings, enabling base-level coordinate conversion. From these, a pseudo-hybrid intermediate genome was constructed by selecting the longer edge between B6 and FVB, thus forming a unified coordinate system encompassing diversity of both strains.

Guppy v6.5.7 in MinKNOW software v23.04.5 was used to call canonical and modified bases (5mC, 5hmC) from raw current signal files fast5/pod5. Reads were then aligned to both references using Minimap2 v2.26 (Li, 2018), retaining only high-confidence alignments (MAPQ > 20). Strain-specific SNPs and alignment scores were used to assign reads to B6 or FVB haplotypes using WhatsHap v2.8 (Patterson et al., 2015). Methylation of each CpG was calculated using modkit v0.1.2 (https://github.com/nanoporetech/modkit). Importantly, all DNAm data were projected onto the pseudo-hybrid coordinate system, minimizing reference bias during haplotype-specific methylation analysis and allowing for a systematic comparison of methylation data from distinct reference coordinate systems.

### Identification of differentially methylated regions

Methylation levels of each CpG for each sample were extracted from bedMethyl files generated by modkit, which contain per-CpG entries detailing genomic coordinates and read counts. Each file included approximately 21 million CpG sites. These datasets were imported into R to create BSseq objects (Hansen et al., 2012) for downstream DMR analysis. To ensure data reliability, unphased datasets retained only CpGs with ≥3× coverage in at least 80% of samples. For phased data, where coverage was inherently lower, CpGs with ≥1× coverage across all samples were included. DMRs were identified in phased data using the bsseq 1.39.4 (Hansen et al., 2012) and unphased data using dmrseq 1.22.1 (Korthauer et al., 2019) R packages. Dmrseq controls the false discovery rate (FDR) whereas bsseq controls the family-wise error rate (FWER). For the phased data smoothing was performed separately for each allele. Strain-specific CpGs were included during smoothing but were removed prior to DMR identification, following previous work (Wulfridge et al., 2019). This strategy enhances statistical power while reducing spurious DMR detection due to strain-specific sequence differences. DMRs were filtered using the following criteria phased analysis: ≥ 3 CpGs per region, > 5% mean methylation difference, and inverse CpG density ≤ 300 (defined as the number of CpGs divided by the region’s genomic width), unphased analysis ≥ 5 CpGs, and inverse CpG density ≤ 120.

### Annotation of differentially methylated regions

We used annotatePeaks.pl program to annotate DMRs by using UCSC mm10 mouse reference fasta and UCSC mm10 refGene gtf files. That program calls a separate program, assignGenomeAnnotation, to efficiently assign DMRs to possible annotations genome wide. By default, in the case that some annotations overlap, HOMER assigns a priority based on the following: TSS (by default defined from -1kb to +100bp), TTS (by default defined from -100 bp to +1kb), CDS Exons, 5’ UTR Exons, 3’ UTR Exons, introns, and intergenic. Intergenic regions were considered either upstream (TSS) or downstream (TTS) of genes.

To annotate DMRs in phased data, a custom pipeline was employed: The UCSC mm10 GTF file was converted to BED format using a Bash script. A Python script mapped coordinates between the mm10 genome and the pseudo-hybrid intermediate genome. This allowed accurate functional annotation of DMRs in a unified coordinate space representing both B6 and FVB haplotypes.

### Enrichment analysis of DMRs in genomic features

Canonical transcripts of protein-coding genes in *Mus musculus* were downloaded from Ensembl using the biomaRt package v2.58.2 (Durinck et al., 2005) and annotated using GenomicFeatures package v1.54.4 (Lawrence et al., 2013) in conjunction with the UCSC genome annotation TxDb.Mmusculus.UCSC.mm10.knownGene v3.10.0. A subset of these genomic features was then selected to include only the canonical protein-coding transcripts. For CpG enrichment, we used the R computing environment to construct a 2×2 contingency table with the number of CpGs for each genomic feature and DMRs. A Fisher’s exact test was used to assess the significance of enrichment.

### DNA methylation overlap across individual datasets

An R-based pipeline was developed to visualize DMRs between mNPCs and Day 6 neurons, incorporating additional conditions including *Dnmt1-*KO and *Kmt2a*-KOs and Day 2 neurons. Unphased DMR coordinates (from mNPC vs. Day 6 comparisons) were used to extract methylation data from the two experiments. CpGs with zero coverage in any sample were excluded. The remaining CpGs were smoothed with the *BSmooth()* function from the bsseq v1.39.4 package. The *normalizeToMatrix()* function from the EnrichedHeatmap v1.32 package was then used to map smoothed average methylation values from each group to a fixed-width matrix centered on the extended DMR intervals (±5kb) with a 100 bp bin size, and heatmaps were generated using the same package.

### PCA plot of genome-wide DNAm

To assess global DNAm variation, the mouse genome was segmented into 1 kb non-overlapping tiles using the *tileGenome()* function from the GenomicRanges R package. Tiles containing more than two CpGs were retained, resulting in approximately 2.7 million genomic tiles. A custom R function was used to generate a PCA plot based on raw genome-wide DNAm levels across samples.

### RNA isolation and sequencing

RNA was isolated as before using TRI Reagent and purified using ZYMO Direct-zol™ RNA Microprep Kit. RNA concentration was measured using Qubit with the Qubit RNA HS Assay Kit (ThermoFisher, Q32852), and RNA integrity was assessed with a Bioanalyzer using Agilent RNA 6000 Nano Kit (Agilent, 5067-1511) according to the manufacturer’s instructions. For *Kmt2a*-KOs and *Dnmt1*-KOs, the two genes were knocked out individually, together with a scrambled control, in 4 biological replicates each, RNA isolated and sent to Novogene Europe for mRNA Illumina sequencing. For wild-type neurodevelopmental cells, 3 biological replicates were isolated for each timepoint, RNA extracted and purified as before and 200ng total RNA from each sample processed using the ONT PCR-cDNA Barcoding kit (ONT, SQK-PCB111.24), according to the manufacturer’s instructions (version v111-revG). Samples were barcoded and pooled at equimolar ratios, then loaded onto 3x R9.4.1 flow-cells (FLO-PRO002) and sequenced on a PromethION 24 instrument.

### RNA sequencing analysis

Illumina short-read RNA-seq (paired-end reads) with 150bp read length on average was generated from *Dnmt1-*KO and *Kmt2a-*KO mNPCs. The libraries were sequenced at approximately 78 million reads per sample. Reads were aligned to the UCSC mm10 reference genome using HISAT2 (Kim et al., 2019). Alignments with > 50 mapping quality were used in featureCounts (Liao et al., 2014) to create a gene-level count matrix. This count matrix was normalized in R using the cpm() function from edgeR (Robinson et al., 2010), and these normalized counts were used for visualization and filtering, whereas raw count data was used for differential expression analysis. Protein coding genes with > 1 Counts Per Million (CPM) in at least one sample were retained for differential expression analysis with DESeq2 (Love et al., 2014), considering genes with an adjusted p-value < 0.05 as differentially expressed. For long-read ONT RNA-seq, libraries were sequenced with an average of ∼10 million reads and a median read length of ∼1.1kb. Reads were aligned to the UCSC mm10 reference genome with Minimap2 (Li, 2018) using *-uf* and *-ax splice* arguments. The featureCounts was utilized to generate the gene-level count matrix with the *-L* argument for long reads. DE analysis followed the same procedure as for short-read data. To ensure reliability, a histogram plot of unadjusted p-values for all genes with > 1 CPM was created in R. To stabilize variance, regularized log transformation (rlog) was applied to raw expression values of retained genes using the DESeq2 package. PCA was performed on the transposed rlog-transformed expression matrix using *prcomp()*, and the first two principal components (PC1 and PC2) were visualized using ggplot2.

### Obtaining promoters of mouse protein-coding genes for NMF analysis

The *promoters()* function of the R package TxDb.Mmusculus.UCSC.mm10.knownGene v3.2 was utilized to identify UCSC mm10 coordinates of promoters. The promoter definition was considered 2kb upstream and 500bp downstream of transcription start site (TSS) of canonical transcript of each gene. The mouse protein coding gene list was downloaded from https://ftp.ncbi.nih.gov/gene/DATA/gene_info.gz.

### Clustering of EM-KOs by NMF based on promoter DNAm abnormalities

We used NMF on smoothed DNAm levels of promoters of protein-coding genes (Cemgil, 2009). CpGs with more than 3x coverage across at least more than 80% of samples were kept in our analysis. Any promoters with no retained CpGs were discarded. The number of latent factors (parameter k) was chosen to be three based on diagnostic plots of NMF and PPI-interactions of EM components. The brunet model in the R package nmf v0.28 was employed in decomposition.

### TF motif enrichment analysis

For identification of TF motif enrichment in DMRs and promoter regions, the Analysis for Motif Enrichment (AME) tool within MEME-suite v5.5.8 (https://meme-suite.org/meme/tools/ame) was used (Bailey et al., 2015). The top 30k most differentially methylated regions from the *Dnmt1*-KO were tested against a background that included 30k random regions from the mm10 genome that were matched in size distribution and CpG density and did not include any of the *Dnmt1*-KO DMRs (**Supplementary Table 15**). For promoter regions of shared DEGs we obtained mouse protein-coding promoters for canonical transcripts as described above, filtered them for the shared DEGs and used all other protein-coding genes as a background. In both cases, we tested against the mouse HOCOMOCO v11 core database and estimated enrichment using a Fisher’s exact test. Odds ratios were calculated using the following formula:

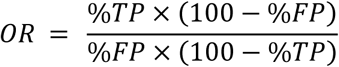

where TP = true positives, or positive hits in the tested sequences, and FP = false positives, or positive hits in the background sequences.

### Inspection of protein-protein interactions using STRING and BioGRID

STRING (Bernas et al., 2017) interactions consisted of three categories: known interactions derived from curated databases and experimental data; predicted interactions based on gene neighborhood, gene fusions, and co-occurrence; and other interactions inferred from text mining, co-expression, and protein homology. We used STRING interactions (confidence score cutoff of > 0.400) between members of the DNAm machinery (Boukas et al., 2019) and the 46 EM-KOs to determine whether the shared impact of EM-KOs on promoter DNAm in **Figure 5A** is associated with these interactions. To identify if TFs with motif enrichments in the *Dnmt1*-KO DMRs or the shared DEGs between *Kmt2a*-KOs and *Dnmt1*-KOs were also known interactors of these proteins, we downloaded all known direct human protein-protein interactions from BioGRID (Oughtred et al., 2021). We filtered the list for known interactors of KMT2A and DNMT1 and compared them with the lists of TFs in R.

### Enrichment analysis and visualization

To test whether the number of overlapping genes between different RNA sequencing datasets was higher than expected, Fisher’s exact tests were performed using the function fisher.test() in R. Shared detected genes were considered background. To test if TF targets or imprinted genes were enriched in DEGs lists, Fisher’s exact tests were performed using the function fisher.test() in R, using detected genes as background. TF target genes were obtained from https://tfbsdb.systemsbiology.net/ (Plaisier et al., 2016) and the TFlink (https://tflink.net/) (Liska et al., 2022) databases. A list of mouse imprinted genes was obtained from the Imprinted Gene Database (https://www.geneimprint.com/site/genes-by-species.Mus+musculus), filtering away genes with “unknown” or “not imprinted” status. To test whether known BioGRID interactors for KMT2A and DNMT1 overlapped significantly, we filtered the human BioGRID list for interactors with either KMT2A or DNMT1. Fisher’s exact test was performed using the fisher.test() function in R, using all interacting partners in the human BioGRID dataset, that are expressed in the two RNA-seq datasets, as universe.

### Gene ontology and gene set enrichment analysis

To estimate enrichment in certain gene ontology (GO) terms in RNA sequencing datasets, the Webgestalt 2024 (Elizarraras et al., 2024) webtool was used, using mouse protein-coding genes as background. The GO term “Biological Process” was evaluated and FDR< 0.05 used to estimate significance. The top 10-15 terms were reported for each analysis. For enrichment of certain GO terms and KEGGs pathways in genes associated with latent factors in the NMF analysis, the EnrichR (Kuleshov et al., 2016) webtool was used. For GSEA of RNA-seq data we ranked the gene lists from *Kmt2a*-KO and *Dnmt1*-KO NPCs using the signed log10 adjusted p-value from the DESeq2 analysis, where the sign was determined by the direction of change. Preranked lists were loaded to the GSEA software (v.4.3.2, BROAD Institute) and enrichment evaluated against the Molecular Signature Database (MSigDB) 2024 Hallmark mouse gene set. Significance was assessed using 1,000 gene set permutations, and results with FDR q < 0.05 were considered significant.

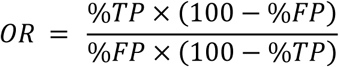

### Data visualization

For scatter plots and volcano plots, the R package ggplot2 was used (Wickham, 2016). For Venn diagrams, the R package VennDiagram was used (Chen & Boutros, 2011). The heatmaps were generated through the *pheatmap()* function of the R package pheatmap. For individual gene expression values from RNA sequencing data, and for quantification results from qPCR, WB and IF experiments, Graphpad Prism (v10.4) was used. Schematic figures were generated using BioRender, and figure assembled into panels using Affinity Designer.

## Data availability

Illumina short-read RNA sequencing data of *Kmt2a* and *Dnmt1* KO mNPCs are deposited in GEO under accession number GSE308011. ONT long-read RNA sequencing data of murine neurodevelopmental samples is deposited in GEO under accession number GSE308012. ONT long-read WGS data of murine neurodevelopmental samples is deposited in GEO under accession number GSE308528. EM KO data is deposited in GEO under two accession numbers GSE308573 and GSE308669. All data will be publicly available upon publication.

## Code availability

Source code for samCRISPR is deposited in https://github.com/kaanokay/samCRISPR. Other source codes are deposited in https://github.com/kaanokay/DNA-methylation-paper.

## Acknowledgements

We would like to thank Elvar Örn Jónsson and Þorgils Árni Hjálmarsson for their dedication and valuable efforts in troubleshooting data storage and analysis issues within the high-performance computing (HPC) cluster at the University of Iceland. We would like to thank Olafur Th. Magnusson for advice regarding ONT sequencing. H.T.B. has a grant from RANNIS (Grant of Excellence, 217988-051) specifically covering this project. J.O. has a grant from Eimskipasjóður Íslands (Doctoral grant, 1238113). K.M. has a grant from RANNIS (Postdoctoral Fellowship, 2410204-052). This work benefitted from equipment from the SAMSNID infrastructure project which brought the PromethION instrument to Iceland, and the IREI infrastructure project which supports the high-performance computer cluster at the University of Iceland. We used ChatGPT (https://chatgpt.com/) to assist in improving scripting efficiency and robustness.

## Author contributions

H.T.B. and K.D.H. conceptualized project and supervised students. H.T.B. received the project funding; J.O., K.M., K.W. S.P., and T.P. performed experiments. K.O., K.M., K.D.H., J.O., S.P, and A.D. analyzed the data. H.T.B., K.D.H., K.M., J.O. and K.O. wrote manuscript. K.A. K.W., S.P., T.P., and A.D. edited manuscript.

## List of supplementary tables

**Supplementary Table 1.** Guide RNA and scrambled guide RNA pairs sequences for each of the genes.

**Supplementary Table 2.** A list of genome-wide significant DMRs from *Dnmt1*-KO, genomic location and presence in regulatory material.

**Supplementary Table 3.** Genomic region enrichment analysis of the *Dnmt1*-KO DMRs.

**Supplementary Table 4.** TF motif enrichment in the top 30K *Dnmt1*-KO DMRs compared to a length and CpG density matched background list.

**Supplementary Table 5.** Differential expression analysis results after *Dnmt1-*KO.

**Supplementary Table 6.** Differential expression analysis results after *Kmt2a-*KO.

**Supplementary Table 7.** Shared DEGs between *Dnmt1*-KO and *Kmt2a*-KO.

**Supplementary Table 8.** TF motif enrichment of promoters of shared DEGs between *Dnmt1-*KO and *Kmt2a*-KO.

**Supplementary Table 9.** Differential expression analysis results after 2 days of differentiation.

**Supplementary Table 10.** Differential expression analysis results after 6 days of differentiation.

**Supplementary Table 11.** Genome-wide DMRs between time points of neurodevelopmental model.

**Supplementary Table 12.** Overlap between *Dnmt1*-KO DMRs and Day6 DMRs.

**Supplementary Table 13.** Haplotype specific DMRs.

**Supplementary Table 14**. Primers used during the current study.

**Supplementary Table 15.** A list of 30k DMRs that were used as background for motif enrichment analysis.

## Supplemental figures

**Supplementary Figure 1:**
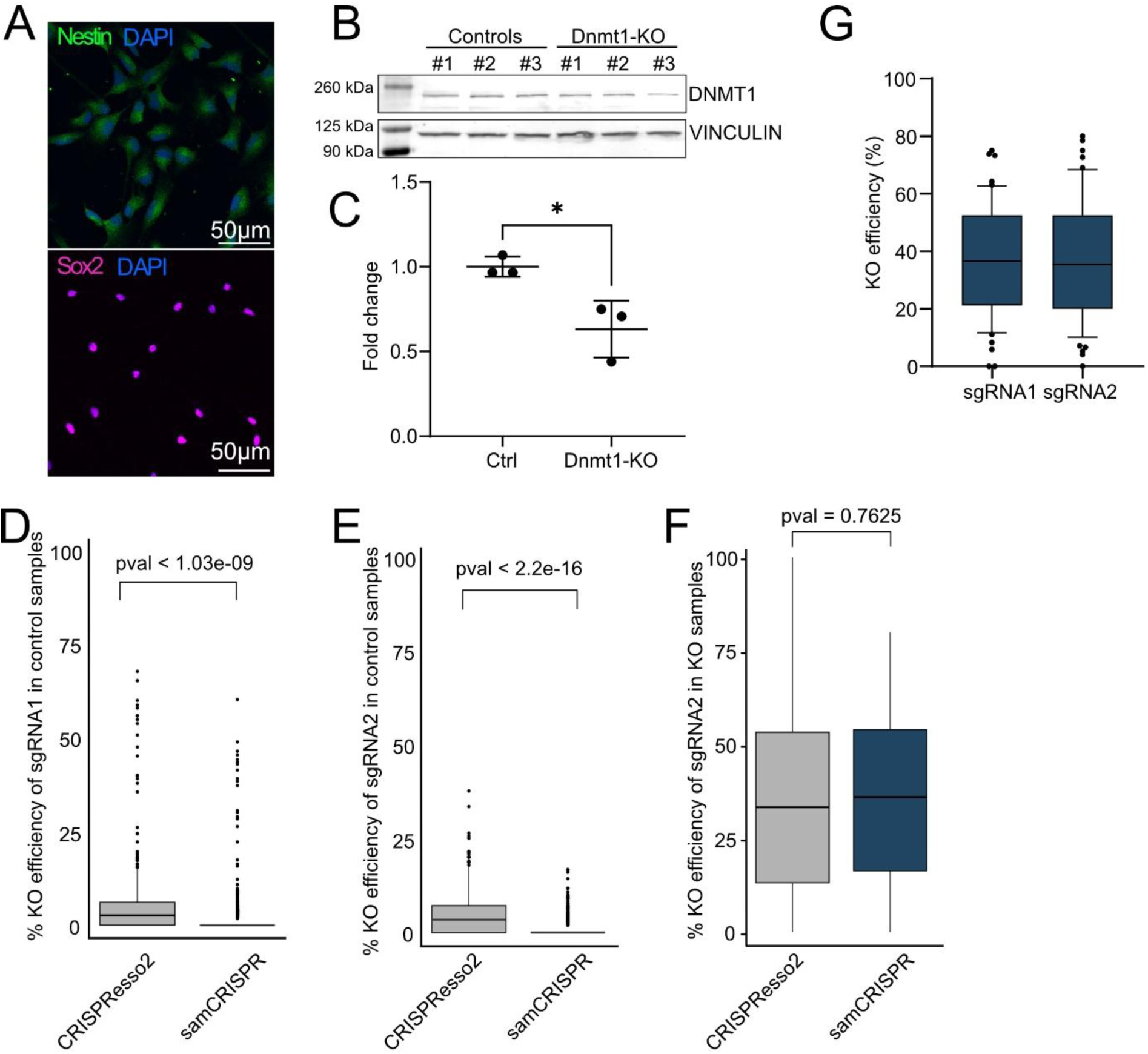
CRISPR-Cas9 KO system evaluation. (A) IF staining of isolated mouse NPCs, staining for Nestin (green, upper) and Sox2 (magenta, lower) and DAPI (blue). (B-C) WB and quantification showing *Dnmt1*-KO and scrambled controls in triplicates. Results in (C) show DNMT1 (upper in B) normalized to VINCULIN (lower in B). A t-test was used for significance. (D) KO efficiency of sgRNA 1 in control samples across methods (two-tailed t-test). (E) KO efficiency of sgRNA 2 in control samples across methods (two tailed t-test). (F) KO efficiency of sgRNA 2 in EM-KOs across methods (two-tailed t-test). (G) KO efficiency of sgRNA 1 and 2 in EM-KOs using *samCRISPR*.

**Supplementary Figure 2:**
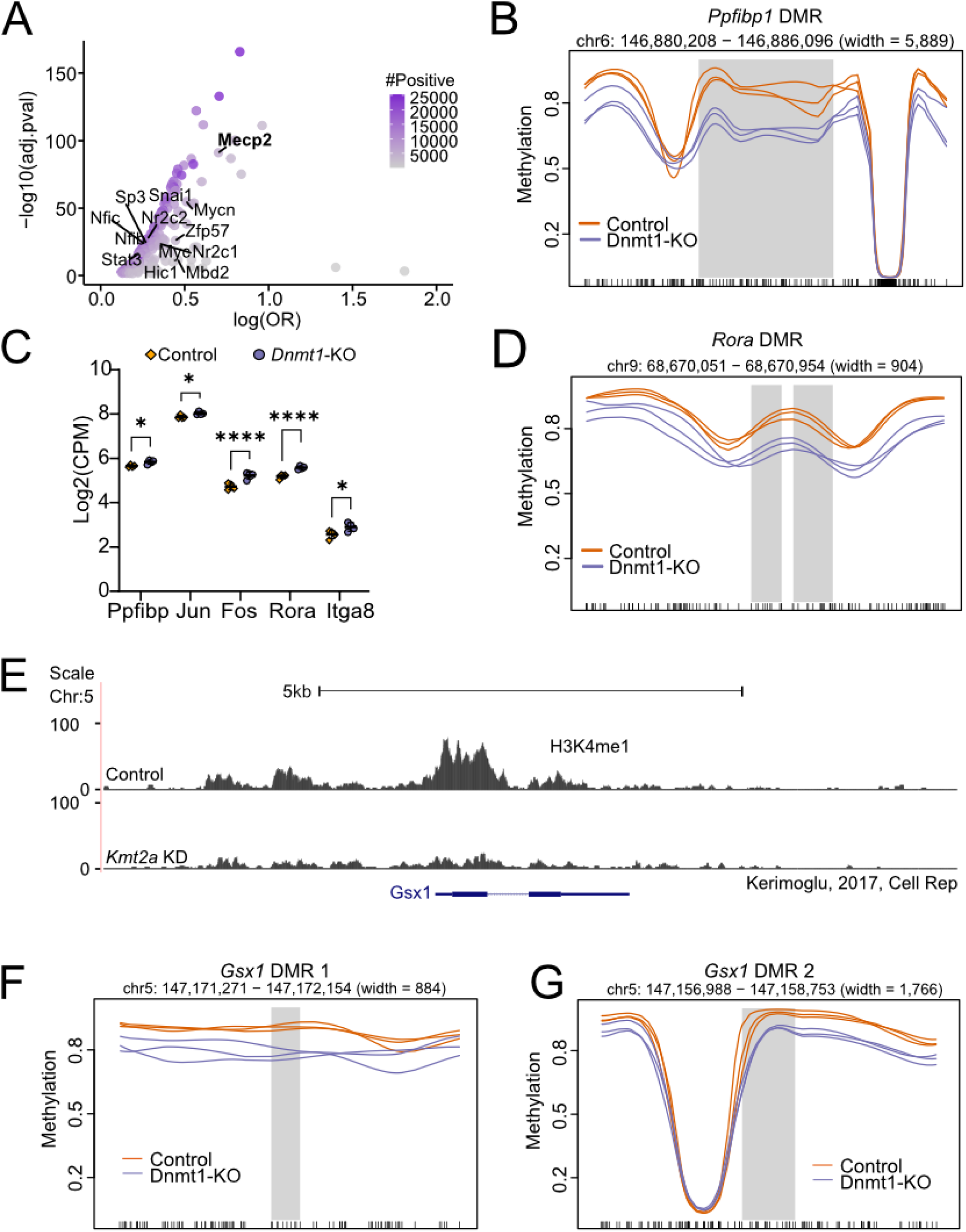
*Dnmt1*-KO and *Kmt2a*-KO DMRs. (A) A scatter plot showing TFs motif enrichment in *Dnmt1*-KO associated DMRs. Known binding partners of DNMT1 from BioGRID are labelled. (B,D) Examples of genome-wide significant DMRs in *Dnmt1*-KO mNPCs (blue) compared to controls (orange), located near the neuronal-related gene *Ppfibp1* (B), and a known DNMT1 target gene *Rora* (D) (Tao et al., 2022). (C) Dot plot showing normalized expression 5 examples of *Dnmt1-*KO DMR-associated DEGs. (E) Publicly available ChIP-seq data (Kerimoglu et al., 2017) showing a loss of H3K4me1 peaks at the *Gsx1* exon in *Kmt2a* knockdown cells compared to control. (F,G) Smoothed methylation plots showing two *Dnmt1-*KO DMRs located in an intergenic region near *Gsx1* gene.

**Supplementary Figure 3:**
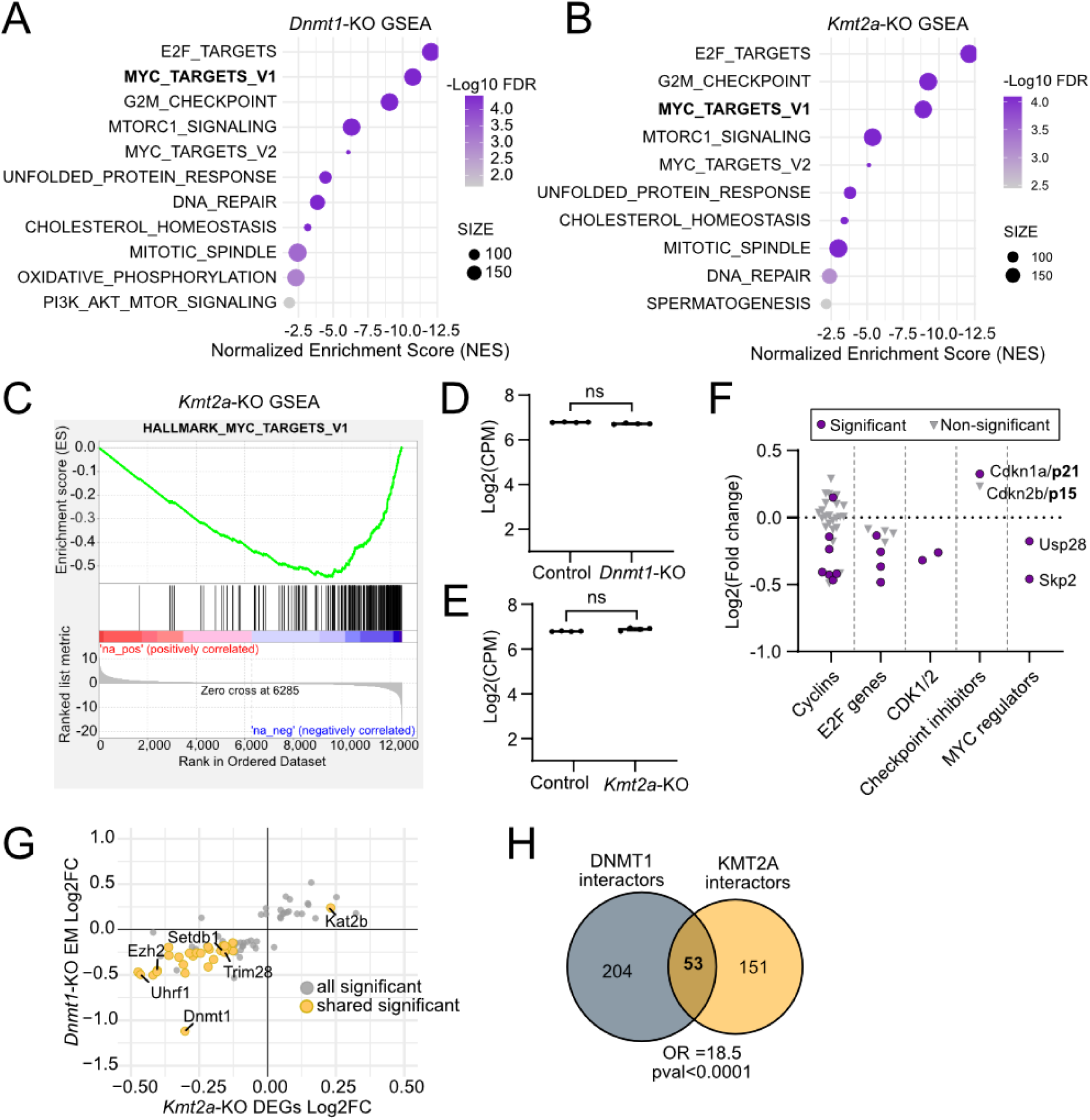
RNA abnormalities in *Kmt2a*-KO and *Dnmt1*-KO DEGs. (A-B) GSEA analysis of (A) *Dnmt1*-KO and (B) *Kmt2a*-KO genes, tested against the MSigDB HALLMARK gene set. The x-axis shows the normalized enrichment score; color denotes FDR adjusted q-values and size indicates the number of genes tested in each category. (C) GSEA plots showing the MYC_targets_V1 Hallmark category for *Kmt2a*-KO genes. (D-E) Log2 normalized *Myc* expression values from the (D) *Dnmt1*-KO and (E) *Kmt2a*-KO RNA-seq data. (F) RNA-seq results showing gene expression changes of MYC targets in *Kmt2a*-KO NPCs compared with controls. Y-axis shows Log2fold changes of genes belonging to the categories on the X-axis. Purple dots show significant genes; gray triangles show non-significant genes. (G) Scatter plot showing log2fold changes of differentially expressed EM genes in *Dnmt1*-KO on y-axis and *Kmt2a*-KO on x-axis. Grey dots show significant EM genes in either one of the datasets, yellow dots show EM genes significant in both datasets. (H) Venn diagram showing the overlap of expressed BioGRID interactors between KMT2A and DNMT1, Fisher’s exact test OR=18.5, p < 0.0001.

**Supplementary Figure 4:**
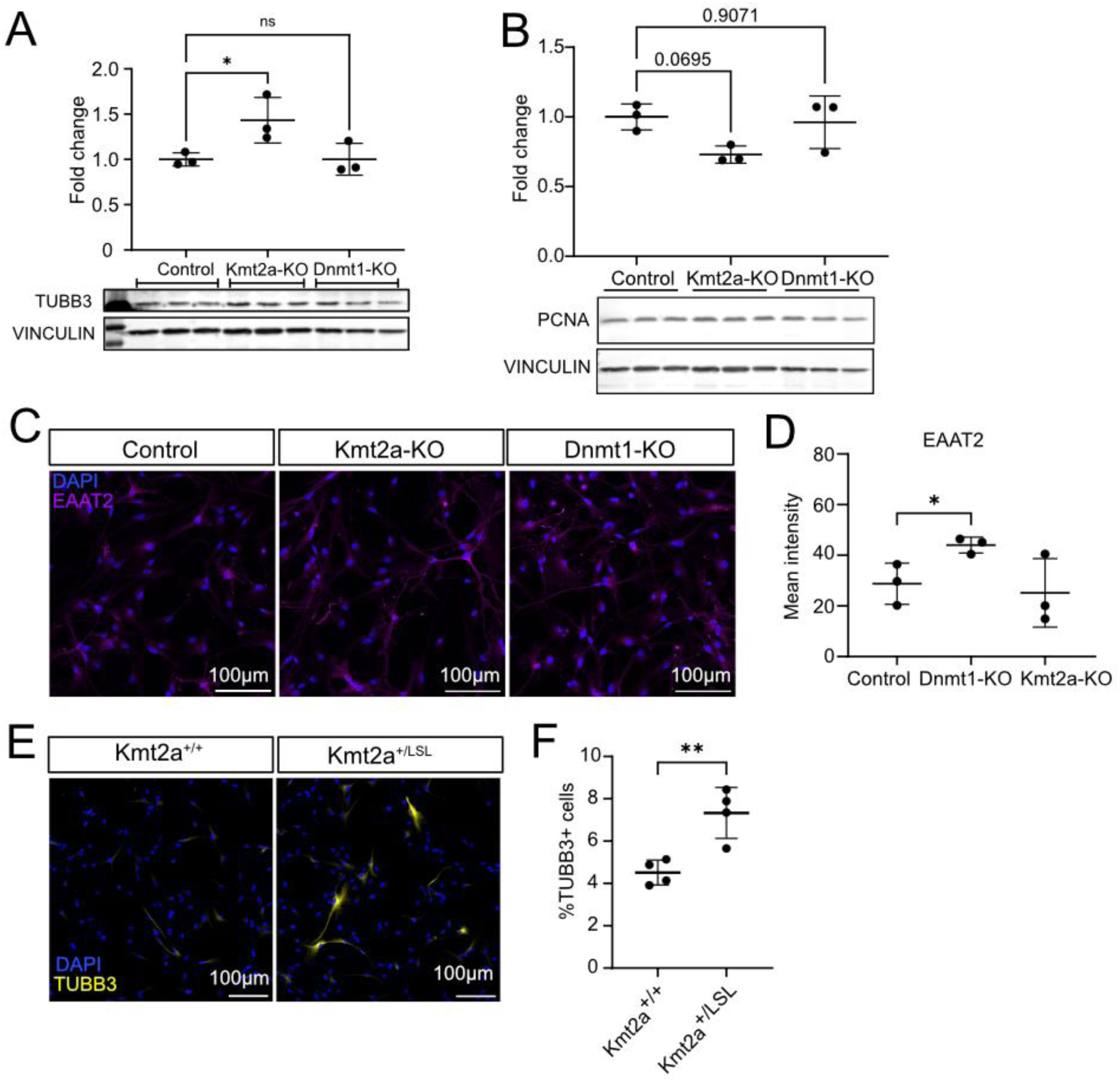
Premature differentiation after *Dnmt1*- and *Kmt2a*-KO. (A) WB analysis of TUBB3 protein after knocking out *Kmt2a* (padj=0.0474) and *Dnmt1* (padj>0.05, ns) with CRISPR/Cas9. A one-way ANOVA with Dunnett’s multiple comparisons test was used to calculate significance. (B) WB analysis of PCNA expressions in *Kmt2a*-KO (ns, p = 0.0695) and *Dnmt1*-KO (ns, p = 0.9071) compared to control mNPC. A one-way ANOVA with Dunnett’s correction was used. (C) Representative IF images of control, *Kmt2a*-KO and *Dnmt1*-KO mNPCs stained for DAPI (blue) and EAAT2 (magenta). Scale bars: 100 µm. (D) Quantification of EAAT2 mean fluorescence intensity in control, *Kmt2a*-KO and *Dnmt1*-KO mNPCs (n = 3, p < 0.05). A one-way Anova with Dunnett’s correction was used.(E) Representative IF images of B6J-*Kmt2a^+/+^* and B6J-*Kmt2a^+/LSL^*P0 mNPCs, stained for DAPI (blue) and TUBB3 (yellow). (F) Quantification of % of TUBB3 positive cells in B6J-*Kmt2a^+/+^* and B6J-*Kmt2a^+/LSL^* P0 mNPCs. A student’s t-test was used. *p < 0.05, **p < 0.01, ns=not significant.

**Supplementary Figure 5:**
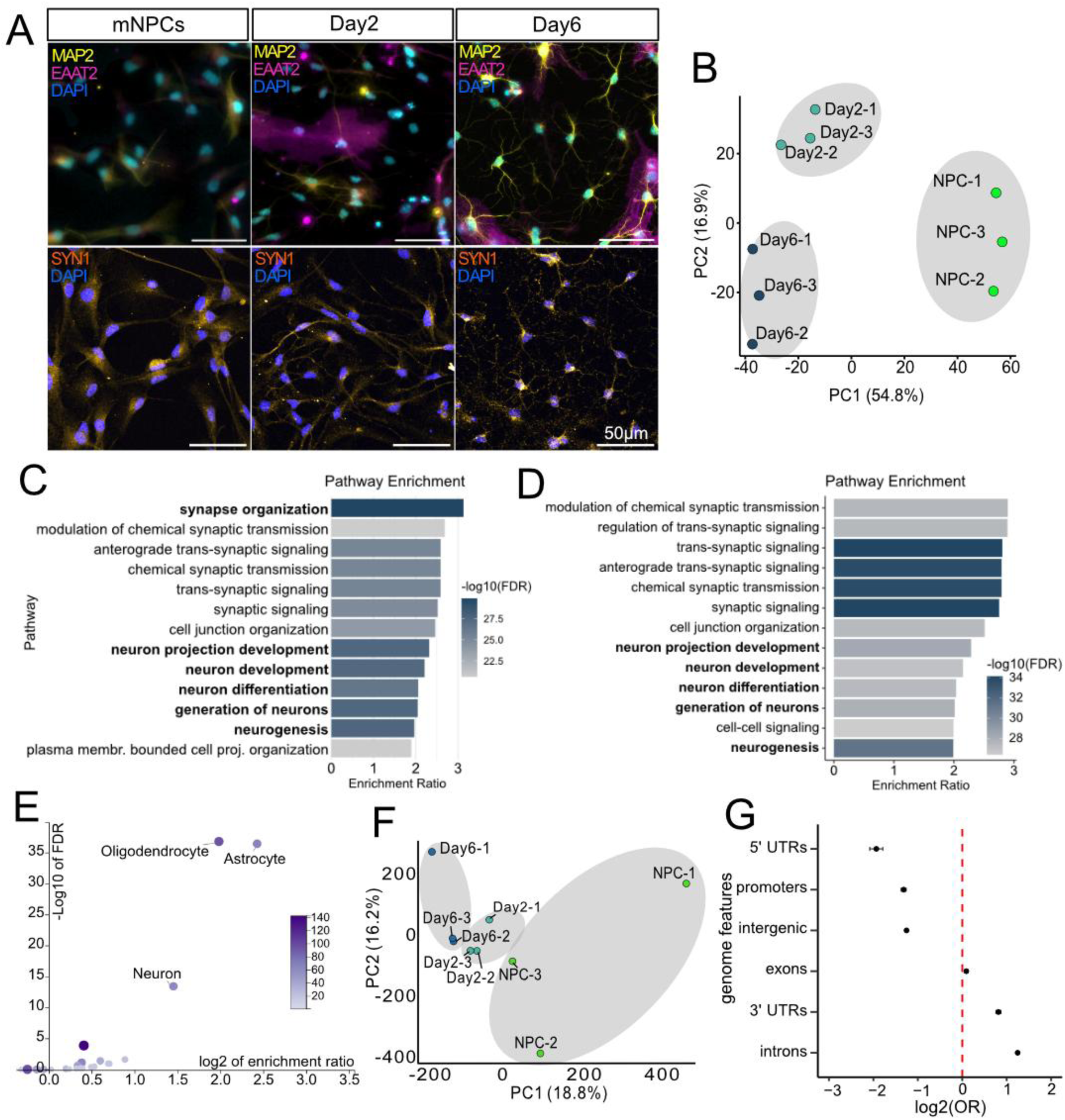
DNA methylation changes in neurodevelopment. (A) IF staining of mNPCs and neuronal cultures at Day2 and Day6 of differentiation. The upper panel shows staining for neurons (MAP2: yellow) and astrocytes (EAAT2: magenta), lower panel shows staining for synapse marker expression (SYN1: orange). DAPI is shown in blue. (B) PCA plot of mNPCs-Day2-Day6 datasets based on ONT RNA sequencing and gene expression analysis. (C-D) GO-term enrichment (Biological Process) analysis showing the top 13 enriched GO terms among DEGs at Day2 (C) and Day6 (D). (E) Gene-set enrichment analysis for cell-types, showing a significant enrichment for all three neuronal lineages in the Day6-DEGs. (F) PCA plot of mNPC-Day2-Day6 samples after ONT DNA sequencing and analysis for differentially methylated regions. (G) Bar graph showing the log2(observed/expected) enrichment ratios for genome locations of Day6-DMRs.

**Supplementary Figure 6.**
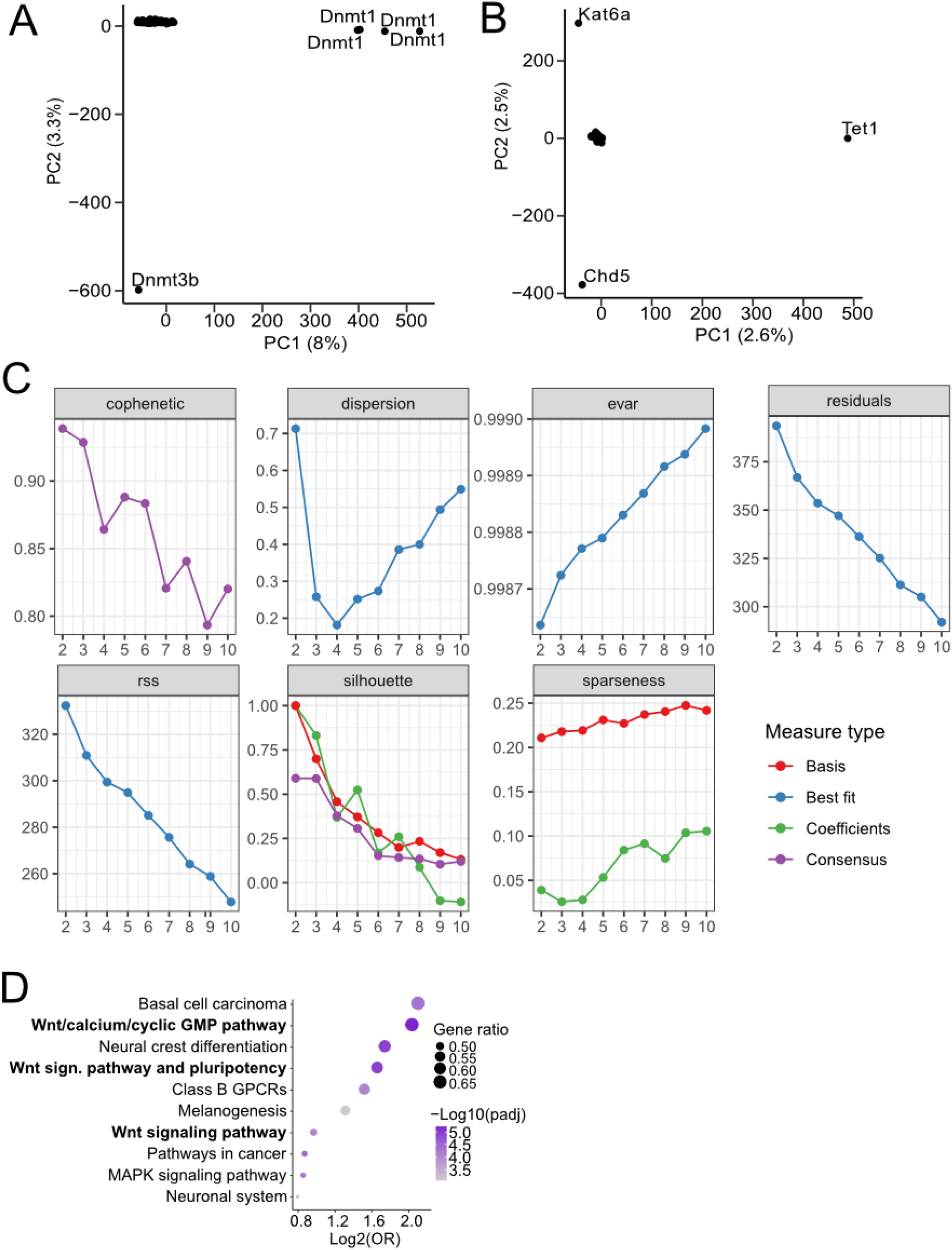
Promoter specific analysis of EM-KO DNAm. (A) PCA plot showing 46 EM-KOs and 8 control samples. (B) PCA plot showing same analysis as in A) but without *Dnmt1* and *Dnmt3b* samples. (C) Diagnostic plots in NMF analysis in decision of number of latent factors (D) GO term analysis (KEGGs pathway) of genes contributing most to latent factor 2.

**Supplementary Figure 7.**
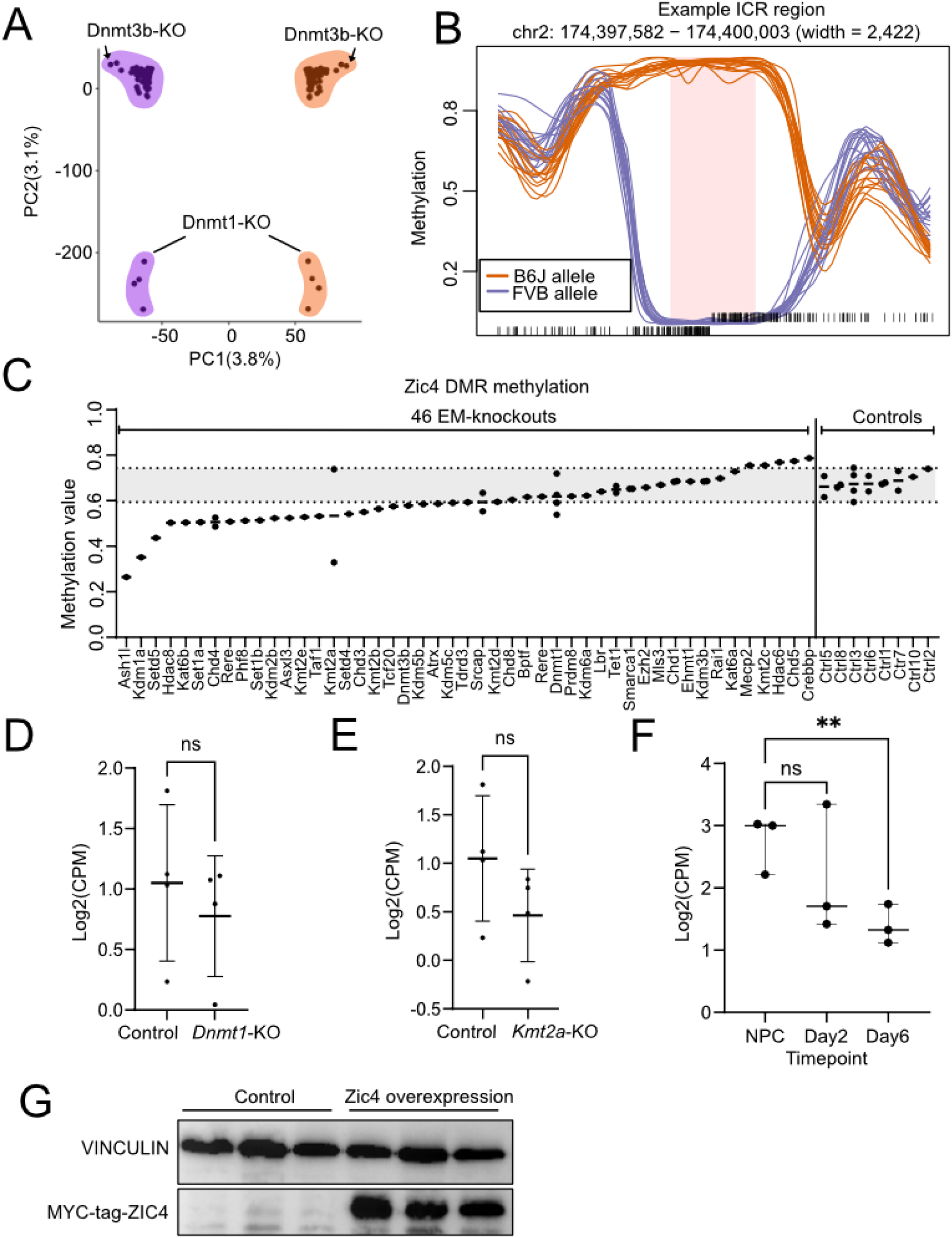
A potential marker gene for EM disruption. (A) A PCA plot showing *Dnmt1*-KO and *Dnmt3b*-KO are two outliers (purple is the B6J haplotype, orange is the FVB haplotype). (B) Smoothed methylation plots showing a haplotype-specific ICR located on chromosome 2. (C) A dot plot showing methylation level at the 3’UTR of *Zic4* locus across 46 EM-KOs compared to 8 controls. Two horizontal dotted lines indicate the minimum and maximum values of the controls. (D-E) Normalized counts showing *Zic4* expression in *Kmt2a*-KO (not significant, p = 0.1846, D) and *Dnmt1*-KO (not significant, p = 0.5, E), compared to controls. (F) Normalized counts showing *Zic4* expression in neurodevelopment, where it is significantly downregulated at Day6 compared with the NPC state (p = 0.00197). (G) A western-blot showing overexpression of Myc-tagged Zic4 and Vinculin as a loading control.

