## Supplemental figures for "Shared molecular consequences of epigenetic machinery disruption in murine neuronal progenitors"

**Supplementary Figure 1**

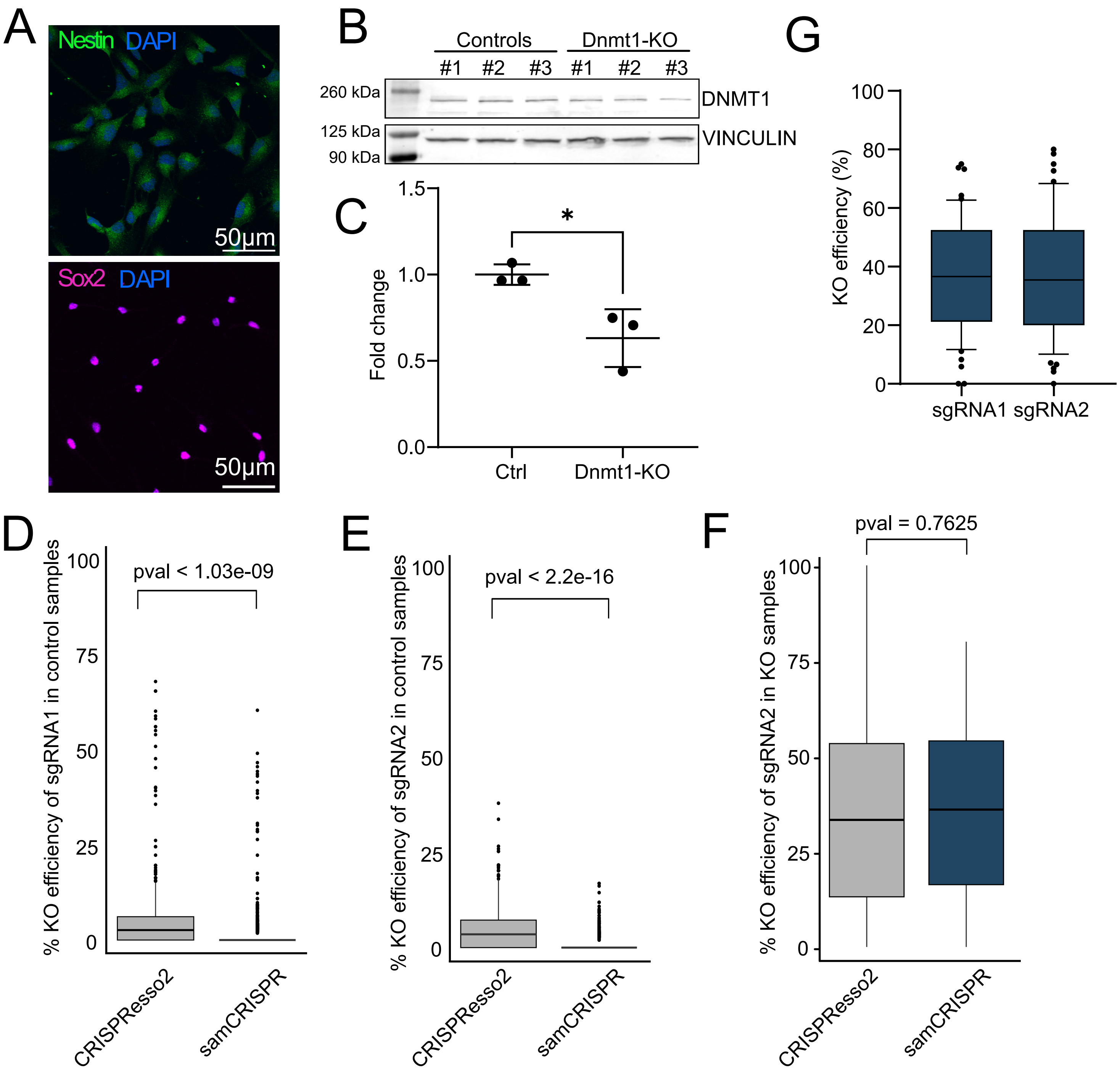

**Supplementary Figure 2**

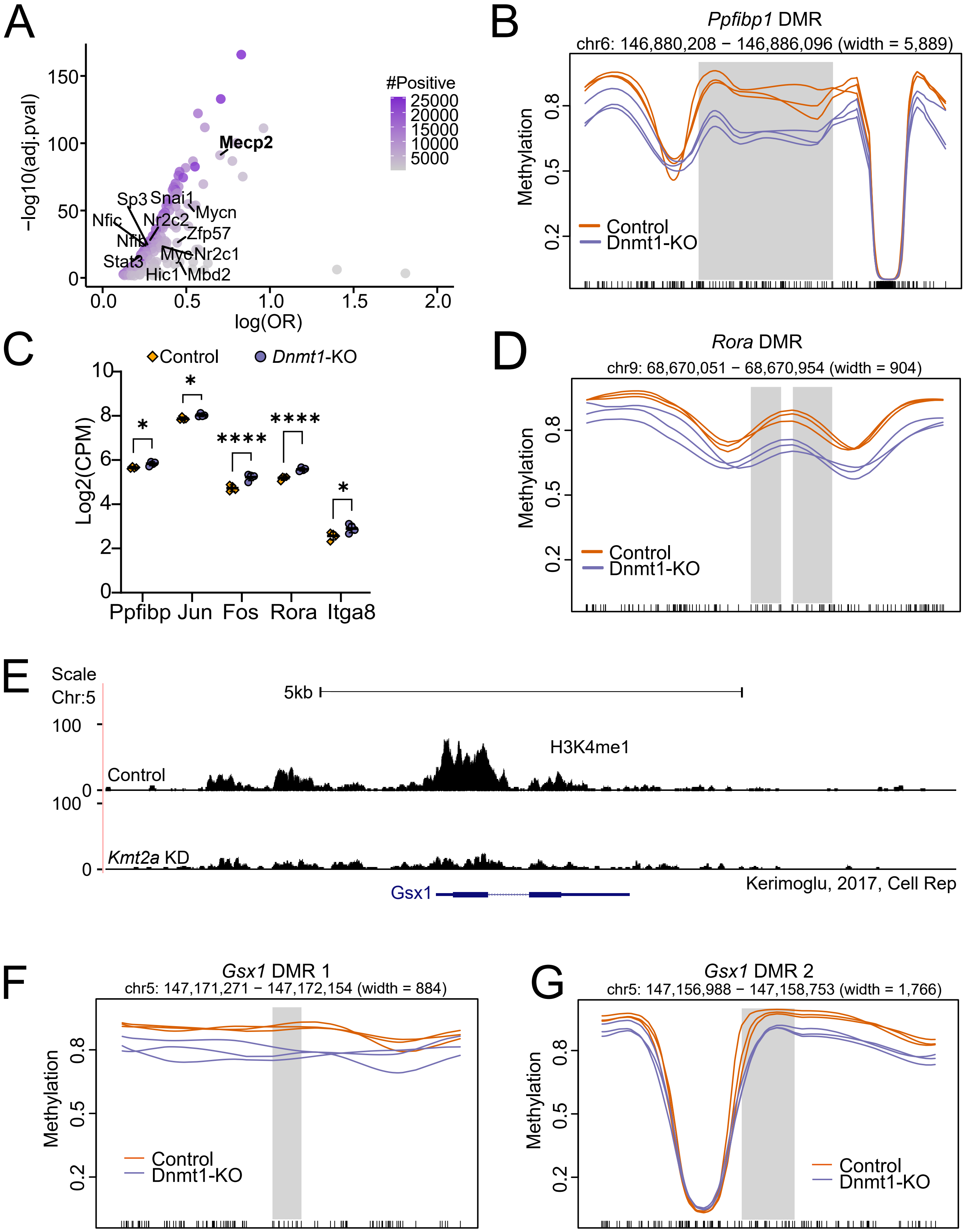

#### Supplementary Figure 3

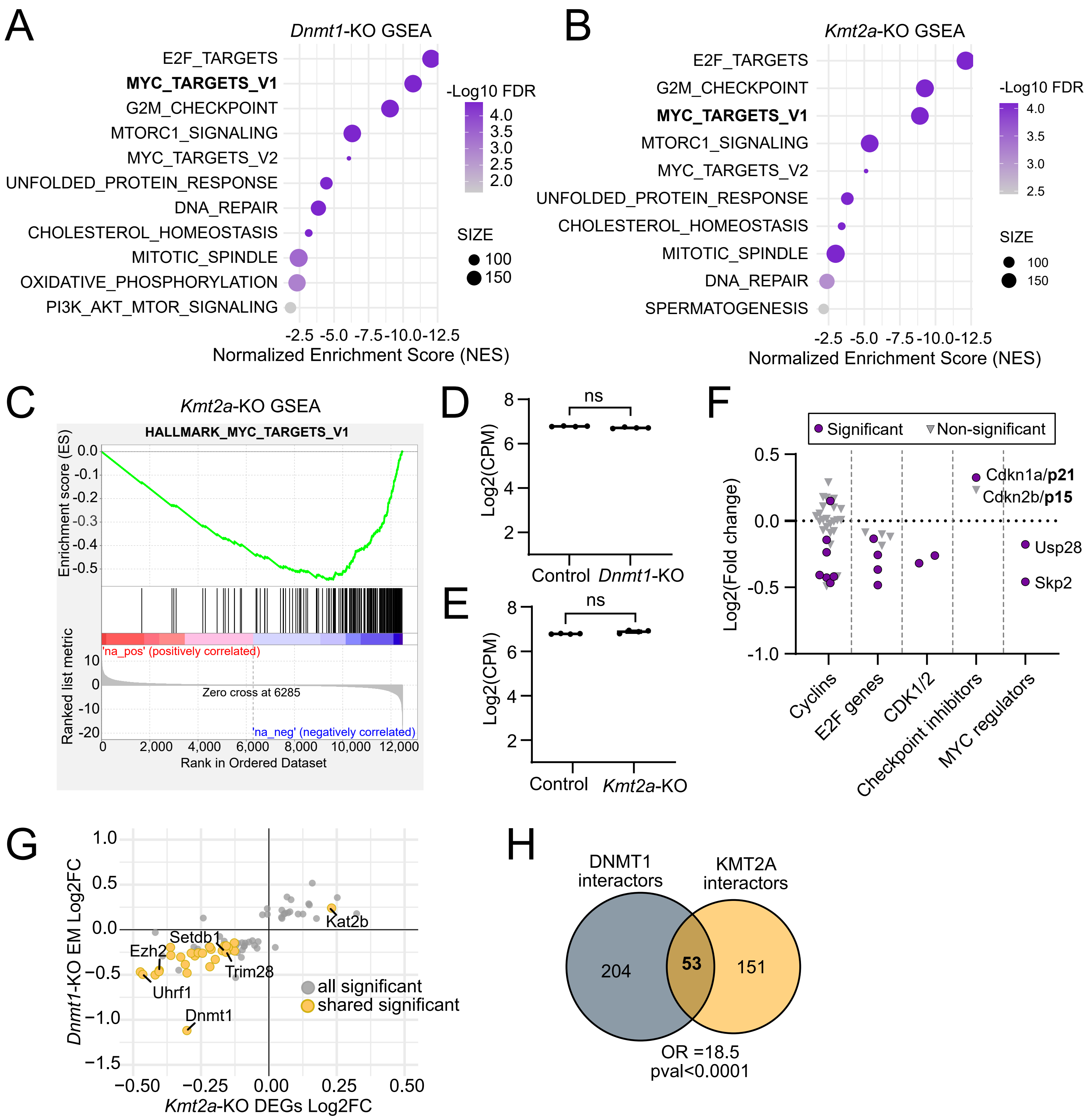

### Supplementary Figure 4

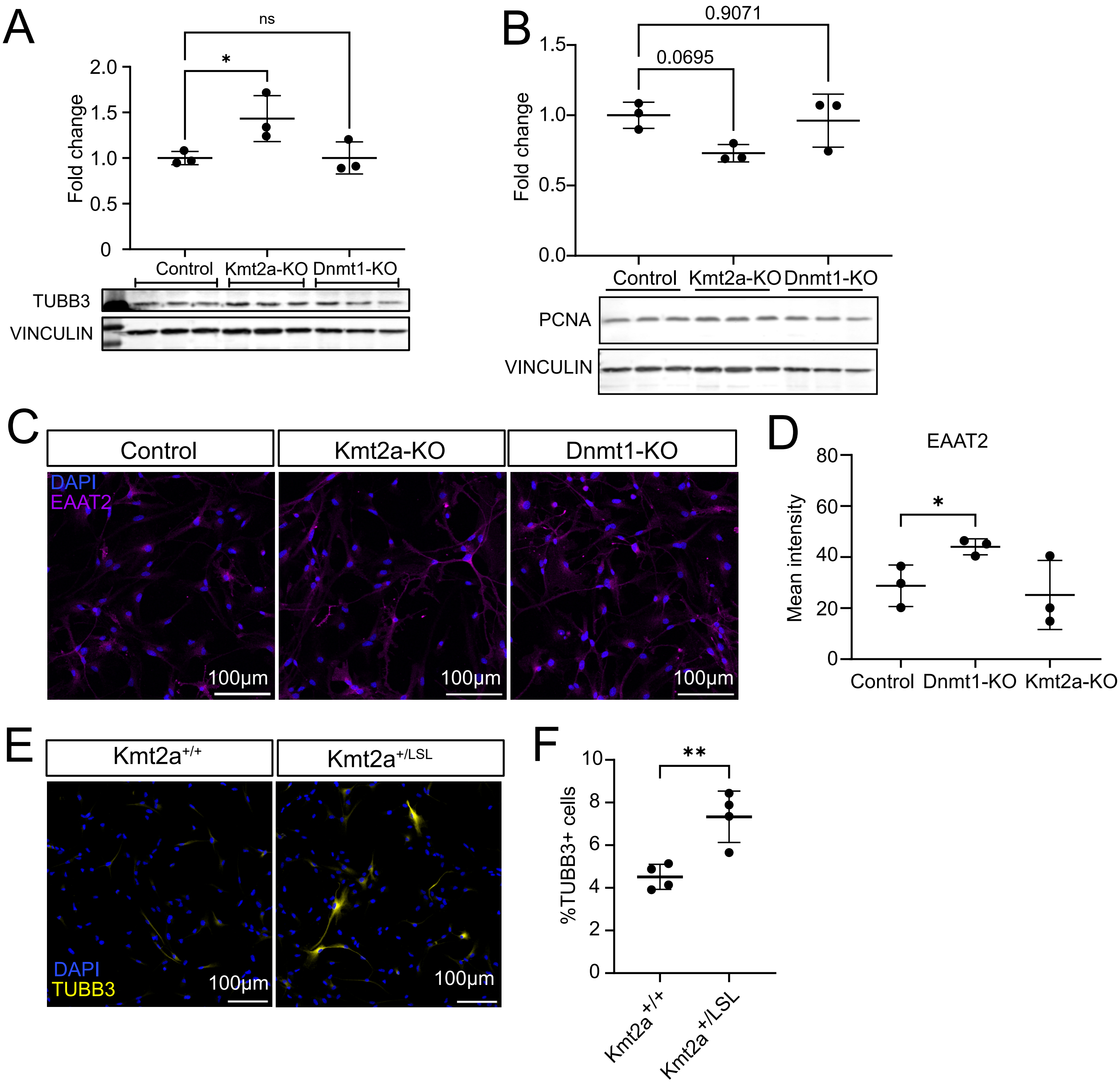

#### Supplementary Figure 5

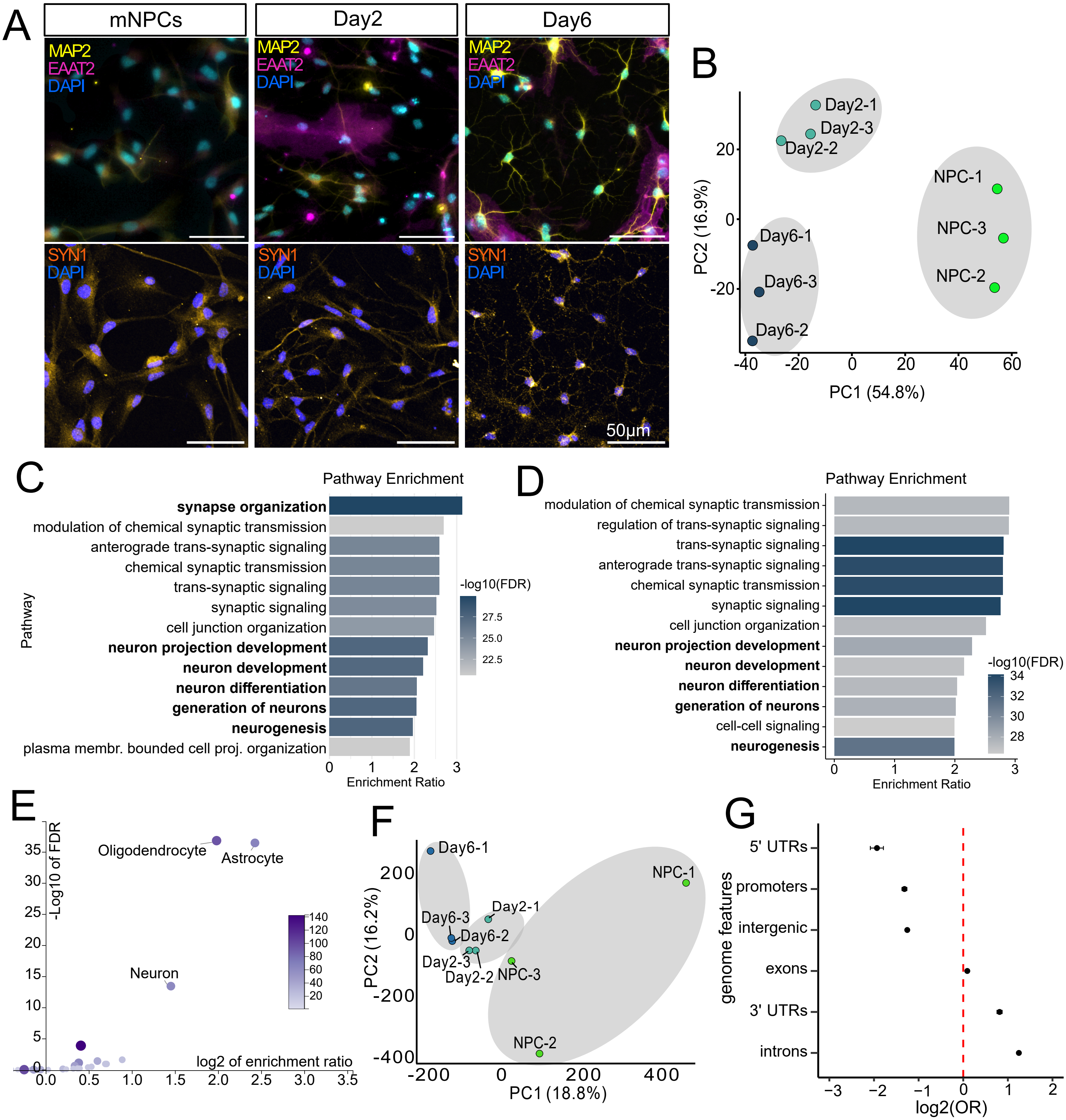

**Supplementary Figure 6**

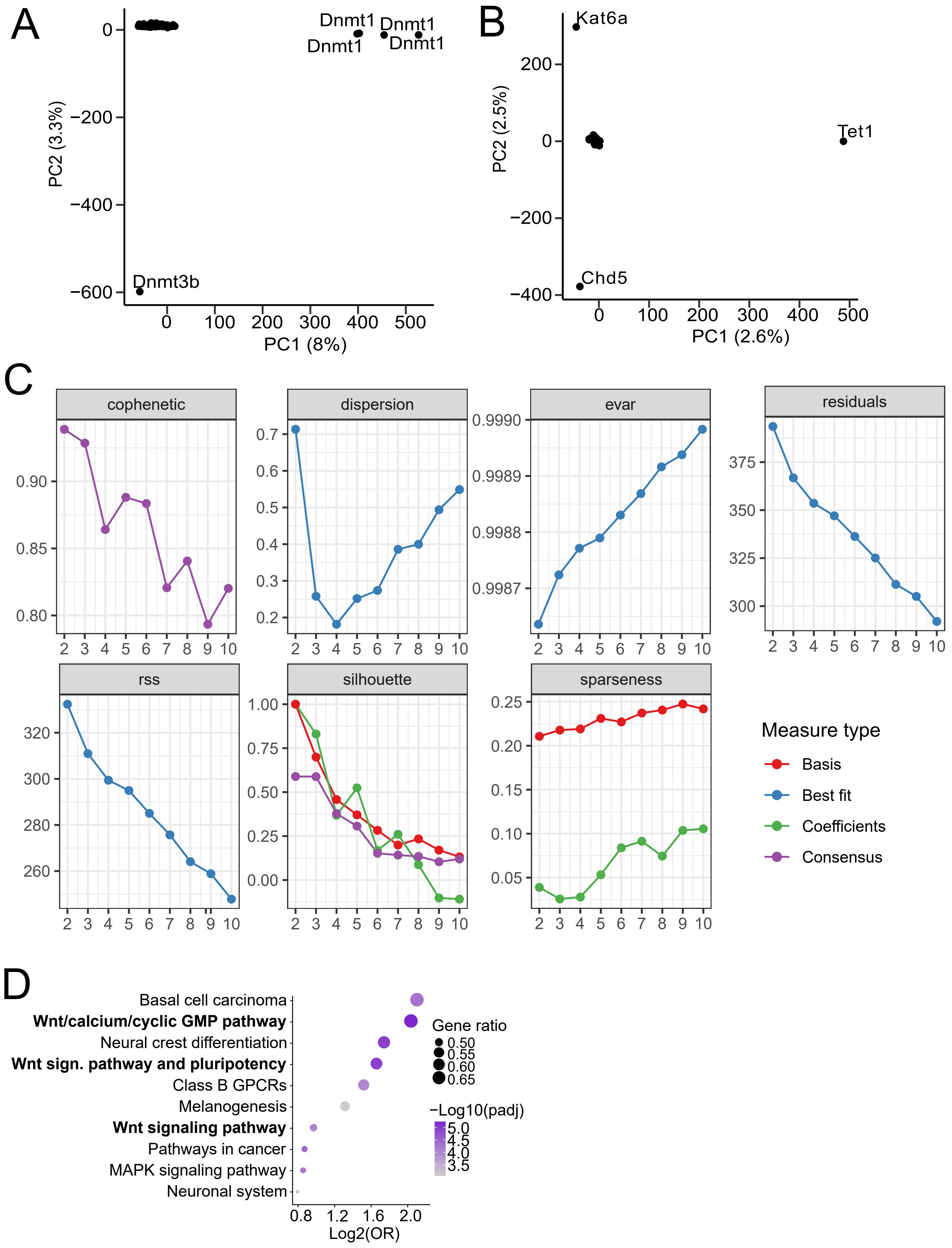

#### Supplementary Figure 7

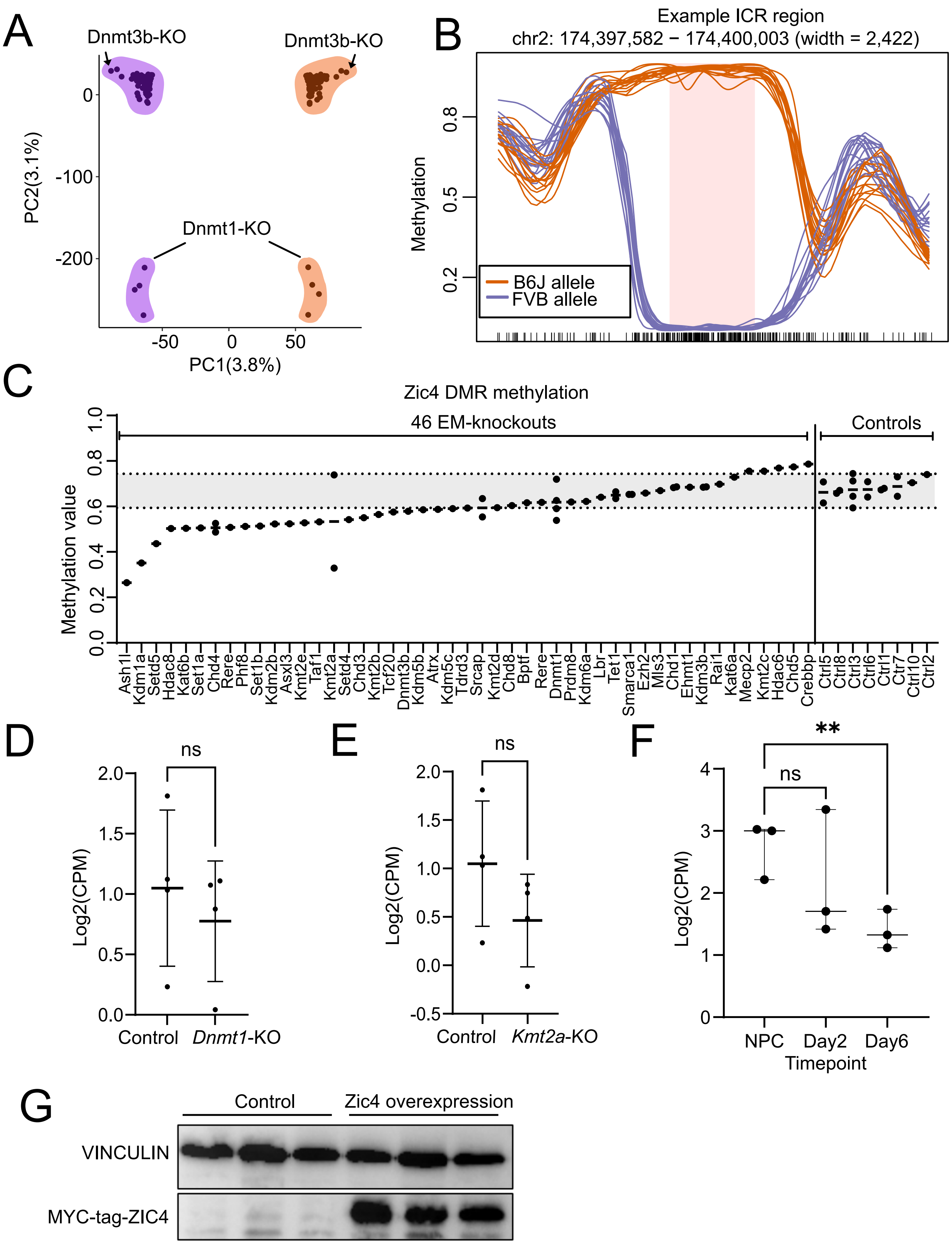
